# An evolutionary optimum amid moderate heritability in prokaryotic cell size

**DOI:** 10.1101/2023.12.18.571935

**Authors:** Henry Secaira-Morocho, Abhinav Chede, Luis Gonzalez-de-Salceda, Ferran Garcia-Pichel, Qiyun Zhu

## Abstract

We investigated the distribution and evolution of prokaryotic cell size based on a compilation of 5380 species. Size spans four orders of magnitude, from 100 nm (*Mycoplasma*) to more than 1 cm (*Thiomargarita*), however most species congregate heavily around the mean. The distribution approximates but is distinct from log-normality. Comparative phylogenetics suggested that size is heritable, yet the phylogenetic signal is moderate, and the degree of heritability is independent of taxonomic scale (i.e. fractal). Evolutionary modeling indicated the presence of an optimal cell size, corresponding to a coccus 0.70 µm in diameter, to which most species gravitate. Analyses of 1361 species with sequenced genomes showed that genomic traits contribute to size evolution moderately and synergistically. In light of our results, scaling theory, and empirical evidence, we discuss potential drivers that may expand or shrink cells around the optimum and propose a stability landscape model for prokaryotic cell size.

## Introduction

While all prokaryotes (bacteria and archaea) come in microscopic sizes, species span several orders of magnitude in cell size, ranging from micrometers up to millimeters ^1–3^. Most are single-cell organisms, although exceptions do exist as some cyanobacteria ^4^, myxobacteria ^5^, and filamentous sulfur bacteria ^6^ are truly multicellular. Cell size holds fundamental significance as it may constrain physiological processes such as the flux of molecules across membranes <u>7</u>, which in turn may determine rates of growth and metabolism ^8^. The “giant” bacterium *Thiomargarita magnifica*, for example,with 1cm-long cells ^9^ exhibits unique adaptations in genomic architecture and cytoplasmic layout to overcome limitations in nutrient acquisition and information processing that arise under the new size regime. In fact, it is often tacitly assumed that the complexity of an organism is dictated by its size ^10^, as exemplified in the expression “higher organisms”.

Cell size is regarded as a multifactorial trait due to its dependance on various genetic and environmental factors ^2,11^. Arguments on the role of environmental factors on cell size tunability range from nutrient availability, with higher nutrient concentrations leading to larger cells ^12,13^, to predator-prey interactions, with smaller cells more likely to escape predation ^14^. While cell size can vary in a single species during the cell cycle and according to external factors ^15^, such variations are small compared to the variation in cell size among species. In fact cell size (along with shape) is perhaps one of the most characteristic and fundamental traits in bacterial systematics, considered to be rather invariant and diacritic for specific descriptions ^16^. Indeed, studies in a few model bacteria show that cell size is tightly regulated through physiological and molecular mechanisms ^15,17,18^. Specifically, the cell division machinery signals a mother cell to divide ^19,20^ so as to produce daughter cells of predetermined cell size depending on the time elapsed between cell division events ^2^. But much of the research on cell size regulation and its variations pertain to a handful of model organisms, and the processes that set preferred size ranges among different species has not been directly addressed, perhaps in part because a comprehensive picture of cell size variations in the microbial world had yet to be compiled. Because cell size generally typifies a bacterial/archaeal species, and species of widely different cell size exist, cell size must change over evolutionary time. However it is unclear whether changes accrue through direct selection on size or through selection on cellular components and physiological traits covarying with it ^21^. Potential covariates considered in the literature include metabolic and growth rates, genome size, nucleoid size, ploidy, ribosomal gene content, number of ATP synthase complexes, and respiratory complexes ^8,22–28^. The relationship between these traits and cell size has been probed using scaling principles across a diverse set of taxa ^29^. The existence of lower and upper limits to cell size based on biophysical and bioenergetic limitations, involving ribosomal content ^28^ or number of respiratory complexes ^25^ has been proposed. While the evolution of organismal size has been analyzed from a macro-evolutionary perspective using models of diffusion over evolutionary time among mammal species ^30,31^, parallel efforts in prokaryotes are missing.

Here we present a large compilation of species-specific cell size data for bacteria and archaea that we used in a systematic analysis of the diversity and evolutionary patterns of this fundamental cell trait. We compiled data on cell length, width, and shape across multiple sources to maximize the biodiversity coverage and identified species-independent properties of cell size employing both coarse-grained and fine-grained approaches. We also assessed the role of evolutionary history on cell size through phylogenetics and fits to evolutionary models and probed the existence of a size optimum and a stability landscape for cell size among prokaryotes, providing a discussion on the potential drivers that may drive cell sizes around the optimum.

## Results

### A large, species-specific cell size dataset for bacteria and archaea

We collected and curated values of cell length, width, and shape for 5380 bacterial and archaeal species from various sources. For each species, we then derived cell volume (*V*) and surface area (*S*) and calculated the ratio of cell volume to surface area (*V/S*) as a measure of linear size that is independent of its cell shape. Aside from basic morphological traits and derived parameters, entries were directly linked whenever possible (1361 species) with publicly available genomic information. **Figure 1** shows the distribution of species in a width/length space. To accommodate the large span of size encompassed, we used logarithmic transformations throughout. A cursory inspection of **Figure 1** shows that the distribution is far from homogeneous in either width or length across the range of possible sizes. Most species congregate in a relatively narrow cell length range (1–10 *μm*) and a much narrower range of cell width (0.3–1.0 *μm*). Some species are clear outliers, including the “giant bacteria” (*Epulopsicium fishelsoni, Thiomargarita namibiensis*, and *T. magnifica*), as well as the smallest bacteria (*Mycoplasma gallisepticum*, and *M. genitalium*). By far, most species are rod-shaped (77.42%). Cocci are less common (15.8%), and spirals or (unicellular) filaments much rarer (3.16% and 0.4%, respectively). This agrees with the common notion that the basic prokaryotic cell shape is a rod ^32^. The distributions of linear sizes of species with different cell shapes, estimated as *V/S* are overlapping but unequal (inset in **Figure 1**), and their medians are statistically different (**Table S1**). Cocci tend to be larger than other cell shapes, with median sizes following the sequence: cocci > bacilli > filaments = spirals.

**Figure 1.**
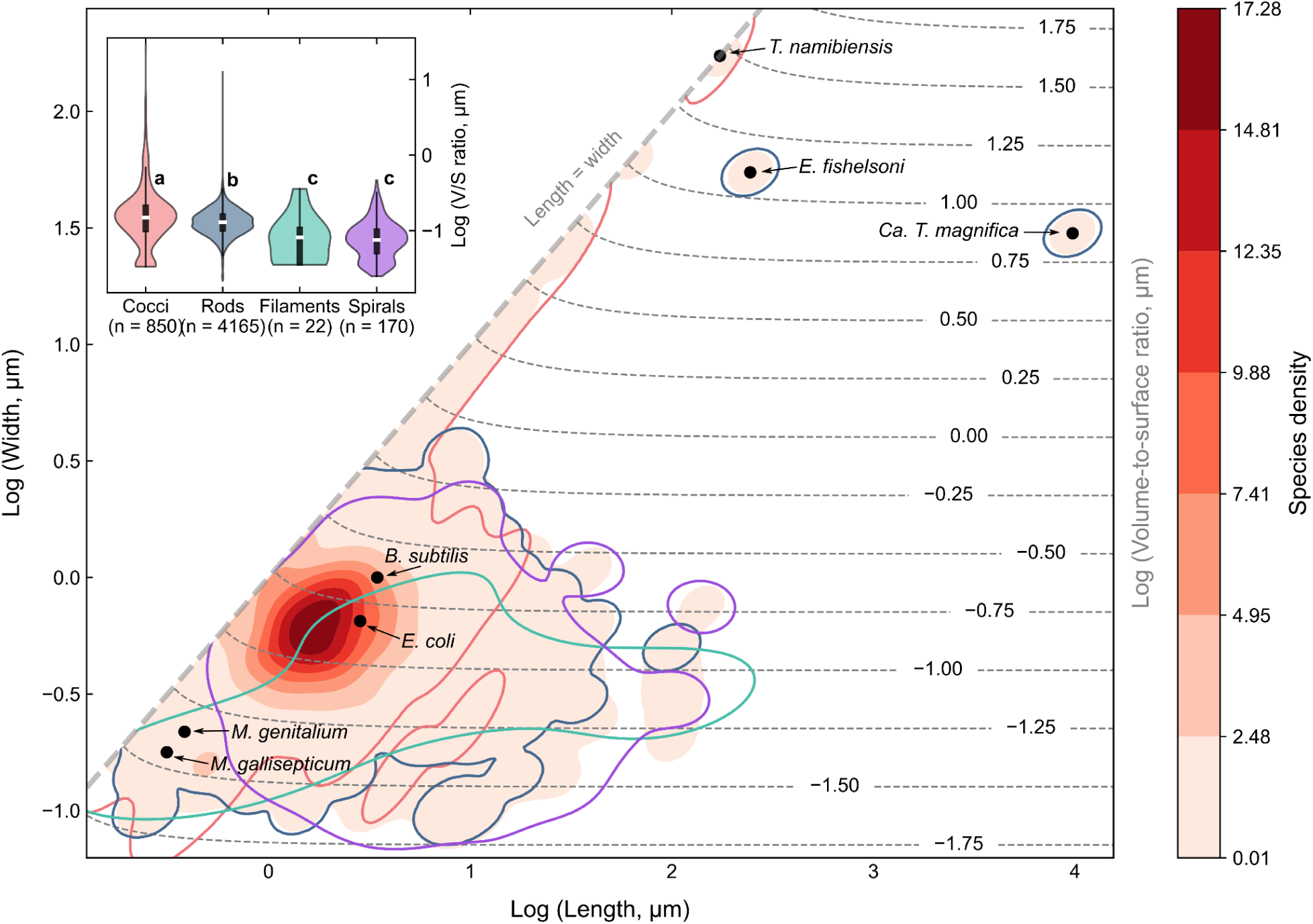
Distribution of cell size among. 5380 **prokaryotic species.** Cell length and width as a heat map of species density. The inset shows violin plot distributions of *V/S* (linear size) binned by cell shapes with median (white line) and interquartile range (black box). Arrows point to large, small, and some model species. Colored lines circumscribe the distributional areas of particular cell shapes, with colors matching the inset legend.

### Cell size distributions

Figure 2A shows the narrow, bell-shaped distribution of log(*V/S*) in our dataset. The mean log*V/S* is ™0.89 *µm*, which corresponds to a linear size *V/S* = 0.13 *µm*, or a coccus with a diameter of 0.78 *µm*. Most species have a linear size quite close to the mean of the distribution. In fact, approximately 95% of species surveyed fall within two standard deviations of the log-distributional mean. While a minor peak on the left-side of the distribution is prominent, we strongly suspect that this is likely an artifact due to rounding of cell length and width values in the source material affecting small-celled bacteria inordinately (as the *V/S* size class enriched corresponds to a reported value of 0.5 *µm* in length). Such artifacts of rounding are also found in surveys of other organisms like insects due to a consensus of using a 1 *mm* boundary for small species ^33^. *V/S* does not conform to a purely log-normal distribution, however, as shown by a Kolmogorov-Smirnov test (*D* = 0.06, *P-value* < 0.0001). Deviation from log-normality is also reported in vertebrate size distributions, but not in insects or phytoplankton ^33–35^. The lack of log normality lies partially in an asymmetrical preference for larger sizes (right skewness value is 0.95, compared to 0 in a normal distribution). Thus, finding prokaryotic species with very large cells, while unlikely, is still much more likely than finding commensurately small species. The right-skewness in logarithmic scaling is a prevalent feature in mammal ^30,36^, lizard ^37,38^, bird, fish, amphibian ^34^, phytoplankton ^35^, and plant ^39^ size surveys. Another reason for the log distribution’s deviation from normality is kurtosis (*K*), or tailedness, which captures the density of outliers on the tails of the distribution ^40^. The distribution is both heavy on outliers and high-peaked, with a higher than log-normally expected abundance close to the mean (leptokurtic with *K* =7.95, where a normal distribution has *K*= 0). **Figure S1** provides a graphical visualization of the differences between the two distributions discussed immediately above. Altogether, these deviations from normality suggest that there might be a range of size that is optimal in most instances and that exceptions in which species have acquired larger cell sizes are more frequent than those involving size reductions throughout evolutionary history, a hypothesis that we attempt to test in the following analyses.

**Figure 2.**
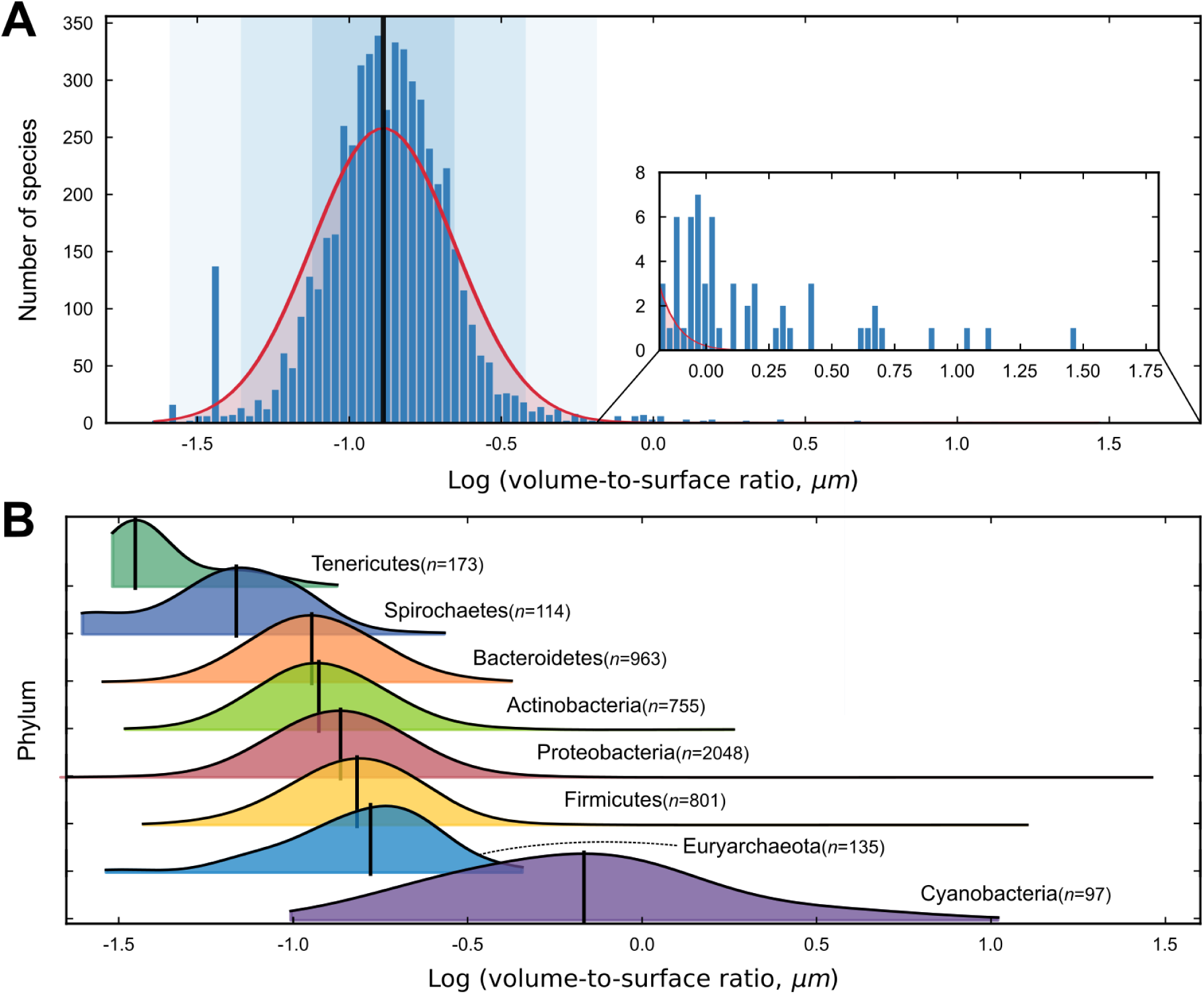
Distribution of volume-to-surface (*V*/*S*) ratios for prokaryotes. **A.** Histogram of log(*V*/*S*) of the 5380 species. The red curve represents the log-normal distribution fit to the data. The black vertical line represents the mean. One, two, and three standard deviations from the mean are shown as a blue background gradient. The mean and standard deviation are log(*μ*) = ™0.89 and log(*σ*) = 0.23, or 0.13 *µm* and 1.71 *µm* on a linear scale. Inset is a close-up for the range of large cells. **B.** Kernel density estimation (KDE) plots of log(*V*/*S*) across phyla. The black lines indicate the medians of the empirical distributions.

One potential reason behind the failure to follow a log-normal distribution is that evolutionary constraints to cell size operate differently across lineages. To test this, we analyzed phylum-specific cell size distributions using kernel density estimation (KDE), which involves placing a Gaussian kernel on each data point. These individual kernels are then combined to create a smooth function that provides an approximation of the probability distribution underlying the data. Figure 2B shows the KDE-derived distribution of log(*V/S*) for those phyla represented in our dataset with at least 100 species. Phyla-specific log(*V/S*) values also approximate bell curves, but with medians that vary markedly across phyla. A Kruskal-Wallis test (*H* = 1133.51, *df* = 7, *P-value* < 0.0001) followed by post hoc Mann-Whitney U tests confirmed that phyla-specific distributions differ significantly (**Table S2**), and each one is significantly different from the rest, highlighting the importance of evolutionary history in the variation of cell size. We found that Tenericutes (a.k.a., Mycoplasmatota) harbor the set of species with the smallest average cell size, whereas Cyanobacteria have the distribution with the largest mean. Most phylum-specific distributions are strictly log-normal according to Kolmogorov-Smirnov tests (**Table S3**), except for Proteobacteria, Firmicutes, and Tenericutes. In the latter, the lack of conformity can be traced back directly to the presence of the artifactual rounding peak (see previous section). In the case of Proteobacteria and Firmicutes, which harbor ‘giant bacteria’, extreme positive skewness contributes to non-conformity (**Figure S2**). Additionally, because phylum-specific distributions, while largely log-normal themselves, have different means and variances, their sum should yield a more complex, compound distribution. This effect may also be behind the lack of log-normality of the entire dataset.

### Cell size diverged through evolution, but only slowly

To further assess the role of evolutionary history on cell size, we asked if the differences among Phyla shown in Figure 2B were the result of a more general, progressive relationship of size diversification with evolutionary distance. To obtain a quasi-continuous measure of evolutionary distance between species, we adopted the Web of Life (WoL) ^41^ reference phylogeny and constructed a phylogenetic tree that includes all species in our dataset (see Methods). We used it to calculate the patristic distance (i.e., the sum of branch lengths connecting two tips in the phylogenetic tree) for every pair of species. For each pair, we also calculated a size difference index (SDI) as the absolute difference in *V/S*, normalized by the maximal difference in the dataset, so that SDI varies between 0 for species of equal size and 1 for species with maximally different sizes. Plotted in Figure 3A are SDIs for pairs of species in the dataset. Both a Spearman’s rank correlation coefficient (*r_s_* = 0.14, *P-value* < 0.0001) and a linear regression (*slope* = 0.02, *P-value* < 0.0001) fitted to all pairs indicate that cell size does diverge during evolution so size differences tend to increase with phylogenetic distance.. The accumulation of variation is more intense, by about an order of magnitude, at patristic distances above 1.88 substitutions per site, which roughly corresponds to taxonomic levels around Phyla. Interestingly, the preferential accumulation of differences in size also occurs at particular taxonomic levels in lizards [family level; ^38^]. Another way of making these patterns obvious is presented in Figure 3B, where we binned a random subset of pairs of species into discrete ranges of patristic distance. Both the progressive increase in distributional means and accrual rates accelerating at large phylogenetic distances are patent. In all, however, the rates of divergence are quite small, with relative SDI medians only increasing by values between 0.01 and 0.08 for either closely or distantly related species, respectively. Altogether, these results show that cell size differences will accrue with evolutionary distance but not fast enough to erase all traces of predictability based on phylogenetic relatedness. Interestingly the standard deviations of the distance-binned distribution also increase with distance implying that the increase in mean SDI through evolution involves an expansion of its distribution, rather than simply a slide towards larger values (**Figure S3**). This is consistent with a random walk or diffusion-like process being at work.

**Figure 3.**
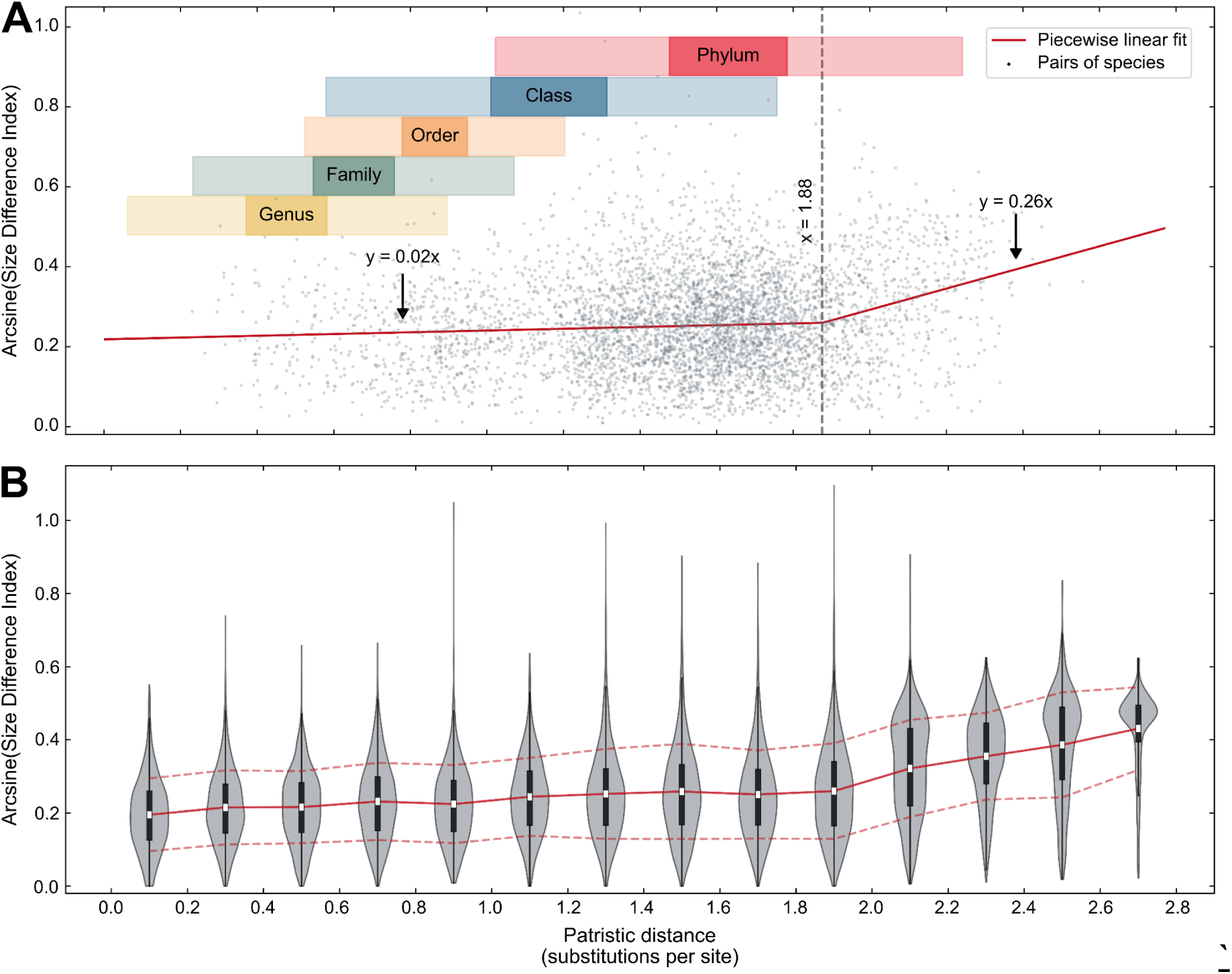
Variations in cell size accrue with evolutionary distance. **A.** Relationship between Size Difference Index (SDI; arcsine-square-root transformed) and patristic distance species pairs. The red line is a piecewise linear fit (*R^2^* = 0.03) while the dashed line indicates the breakpoint in the slope. Displayed as gray points is a randomly selected subset of (0.003%) all pairs of species considered, for the sake of clarity. Colored bars indicate the approximate patristic distance range corresponding to various taxonomic units. **B**. Violin plots of arcsine-square-root transformed SDI, binned by patristic distance ranges (*n* = 1000 pairs per bin). White dots indicate means and black boxes the interquartile ranges. The red line shows the trend in medians, and dashed lines bound the standard deviations.

### Heritability of prokaryotic cell size

One can formally quantify the dependence of a trait on evolutionary history using evolutionary models. In the Brownian Motion (BM) model, the trait under consideration changes with time according to a random walk (its probability density function is normal) that results from the addition of small stochastic and directionless forces affecting the trait ^42^. The model Pagel’s *λ*, which varies between 0 and 1 and is also known as “phylogenetic signal”, quantifies the goodness of fit of a dataset to the BM model. If *λ* = 1, variability in the trait is fully explained by evolutionary history. If *λ* < 1, other factors unrelated to evolutionary history must have an impact. If *λ* = 0, then evolutionary history plays no role. We found that *V/S* had a significant phylogenetic signal (*λ* = 0.83; *P-value* = 0.00). Because it has been suggested that values of Pagel’s *λ* can be sensitive to the number of species used in estimation ^43^, we evaluated the robustness of our estimate by running saturation curves with increasing sampling effort, finding that taxon bias does not have a strong effect on the estimate (**Figure S4**). As an alternative, the parameter, Blomberg’s *K*, which ranges from 0 to infinity, also quantifies the strength of the phylogenetic signal under a BM model, as the ratio between the variance among clades to the variance within clades. A high ratio speaks for a high phylogenetic signal and vice-versa; a ratio of 1 is the expectation of the BM model ^44^. We found that in our dataset, *K* = 0.38 (*P-value* = 0.01), supporting the presence of a certain, albeit not strong, phylogenetic signal. For a cross-trait comparison, we also measured the phylogenetic signal in the same dataset of several genomic properties, obtained through metadata of the WoL project (**Table 1**). Clearly, the inheritability of cell size, while detectable, is much weaker than that of basic genomic properties like genome size or GC content, except for the 16S rRNA gene copy number per genome, which is less inheritable than cell size. Since cell size is likely a complex, integrative trait, the comparison with simple genomic properties may be regarded as improper. And yet the phylogenetic signal of cell size is still weaker than that of another complex trait, the average optimal growth temperature. Overall, we show that evolutionary history plays a role in explaining the variation of cell size because cell size displays a certain degree of conservation and inheritability over broad phylogenetic scales, but inheritance controls size variation more weakly than it does in other traits over evolutionary time.

**Table 1.** Phylogenetic signal of various traits (*λ*: value of Pagel’s *λ*; *σ*^2^: variance; *K*: value of Blomberg’s *K*; *P: P*-value; ΔENC’: effective number of codons; *T*_opt:_ average optimal growth temperature).

| Trait | Number of species | Pagel's $\lambda$ | | | | Blomberg's $K$ | |
| --- | --- | --- | --- | --- | --- | --- | --- |
| | | $\lambda$ | $\sigma^2$ | $P^a$ | Lnlik | $K$ | $P^b$ |
| Log volume-to-surface ratio | 1361 | 0.76 | 0.08 | < 0.0001 | 359.73 | 0.32 | < 0.05 |
| Log genome size | 1361 | 0.99 | 0.05 | < 0.0001 | 1199.05 | 1.28 | < 0.05 |
| Log number of proteins | 1361 | 0.98 | 0.04 | < 0.0001 | 1216.92 | 1.06 | < 0.05 |
| Arcsine of GC content | 1361 | 1.00 | 0.01 | < 0.0001 | 2108.80 | 1.64 | < 0.05 |
| Coding density | 1361 | 0.95 | 0.005 | < 0.0001 | 2597.30 | 0.53 | < 0.05 |
| 16S rRNA copies | 1361 | 0.21 | 4.09 | < 0.0001 | -2764.02 | 0.13 | < 0.05 |
| $\Delta\text{ENC}'$ | 1361 | 1.00 | 0.01 | < 0.0001 | 2180.82 | 1.06 | < 0.05 |
| $T_{\text{opt}}$ | 3431 | 0.99 | 171.13 | < 0.0001 | -11051.71 | 3.90 | < 0.01 |

<sup>a</sup> $P$ -value for the null hypothesis (no signal; $\lambda = 0$ ).
<sup>b</sup> $P$ -value for the null hypothesis (no signal; $K = 0$ ).

### Cell size phylogenetic signal is independent of taxonomic range

The inheritability of cell size, however, need not be a constant through evolutionary history. To test this, we assessed variations in Pagel’s *λ* across taxa at various levels of resolution (i.e., phylum, class, order, family, genus), as defined in our overall phylogenetic tree (see Methods). Only taxa containing more than 50 species were analyzed to ensure that our estimates were robust (**Figure S3**). We present the results of these analyses as a hierarchical network in Figure 4, with a full statistical analysis in **Table S4**. We could demonstrate that phylogenetic signal varies among clades as far as it is possible to vary, from 0 to 1, but also that there is no clear trend between phylogenetic signal and level of taxonomic resolution. In fact, the variance of Pagel’s *λ* across phyla is as high as its variability across classes, orders, and so on (Levene’s test: *W* = 0.32, *P-value* = 0.81). Thus, because phylogenetic signal can be found at any taxonomic scale and its variability is scale-independent, one could characterize it as fractal.

**Figure 4.**
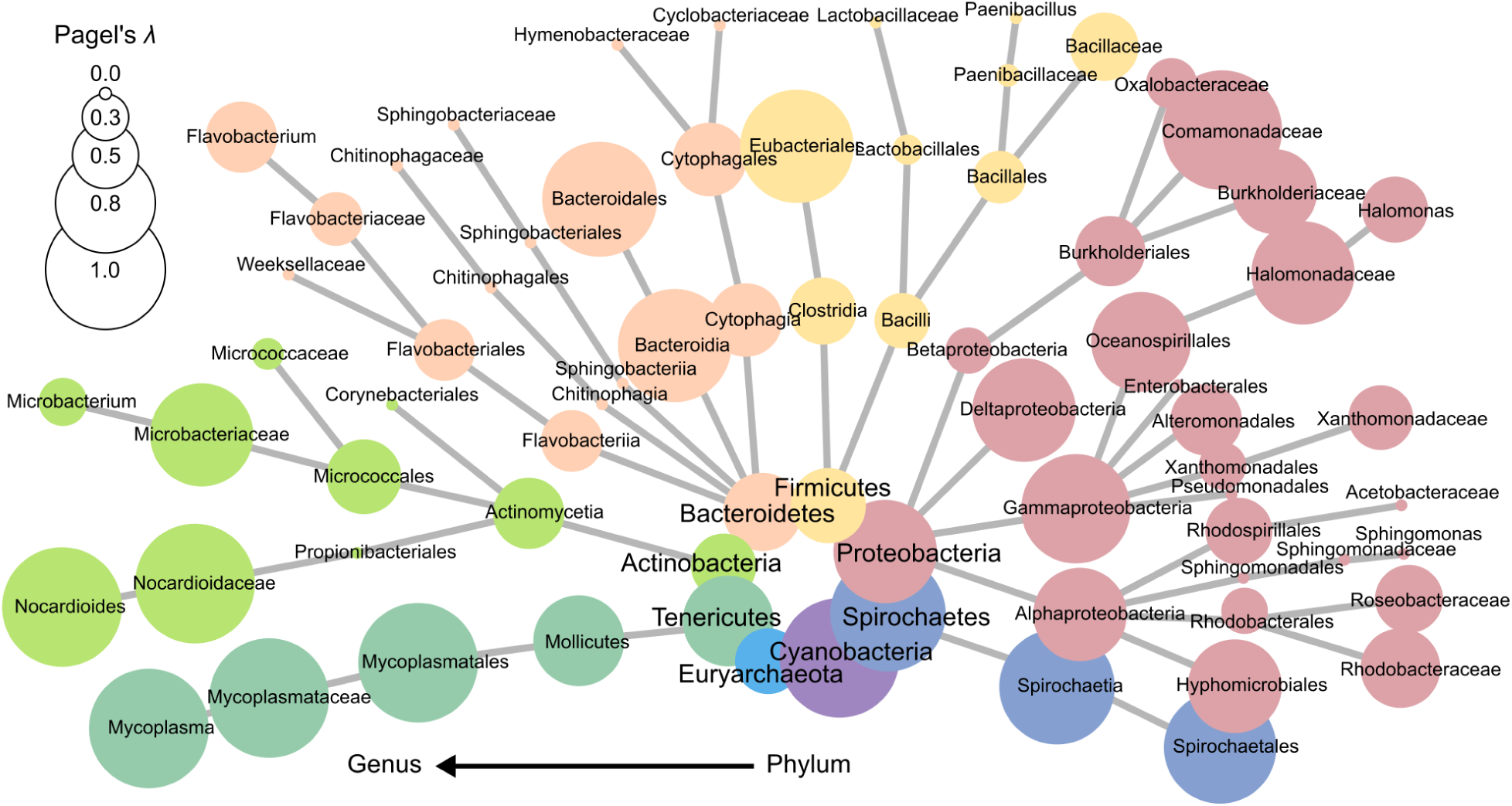
The strength of phylogenetic signal is independent of taxonomic scale. Hierarchical network showing Pagel’s *λ* (as the size of the nodes) within taxonomic groups with more than 50 species. Nodes represent taxonomic groups (color coded according to phylum) and finer taxonomic resolution is placed increasingly outwards.

### Evolutionary models speak for the existence of an optimal cell size in prokaryotes

To identify a plausible model of evolutionary descent in cell size among prokaryotes, we adopted a hypothesis-testing framework, evaluating comparatively four evolutionary models: Brownian Motion as explained above ^42^, Early-Burst (EB), in which the rate of evolution decreases through time ^45^, White-noise, in which phylogeny is irrelevant, and Ornstein-Uhlenbeck (OU), in which the trait returns towards an optimal value ^46^, fitting our data to each of the models, and weighting goodness of fits using Akaike weights. By far the best fit was under the OU model (**Table 2**) which yielded the highest log-likelihood and smallest (most negative) Akaike Information Criterion. The combined weight of the evidence for the latter criteria is summarized in the Akaike weights ^47^, which leaves few doubts about the strength of the OU model. This model, in turn, suggests that there is an optimum value (*θ*) for cell size (log(*θ*) = ™0.93), whose value is close to the log mean (log(*μ*) = ™0.89) of the *V/S* distribution from Figure 2. *θ* measures the long-term mean around which traits evolve, so both the proximity and inequality between *θ* and *μ* were not unexpected ^48^. The strength of attraction towards the optimum in the OU model is gauged by *α*, which takes positive values equal or larger than 0, where *α* = 0 denotes a purely Brownian type of evolution with no attraction towards an optimum. When *α* is a large positive number, then the imprint of trait variation through time and the phylogenetic signal are lost. Because *α* scales with tree height, interpretation is easier if the tree is rescaled to a maximal distance of one, as we did, and gauging by −log(*α*) ^48^. A value of −log(*α*) close to 4 denotes Brownian evolution, whereas values close to ™4 indicate a strong tendency to return toward θ. In our case −log(*α*) = ™1.09, which indicates a clear tendency to return towards the optimum that is moderate enough to allow a phylogenetic signal to be detectable. Analyses performed without rescaling the phylogenetic tree also show the OU model as the best fit (**Table S5**).

**Table 2.**
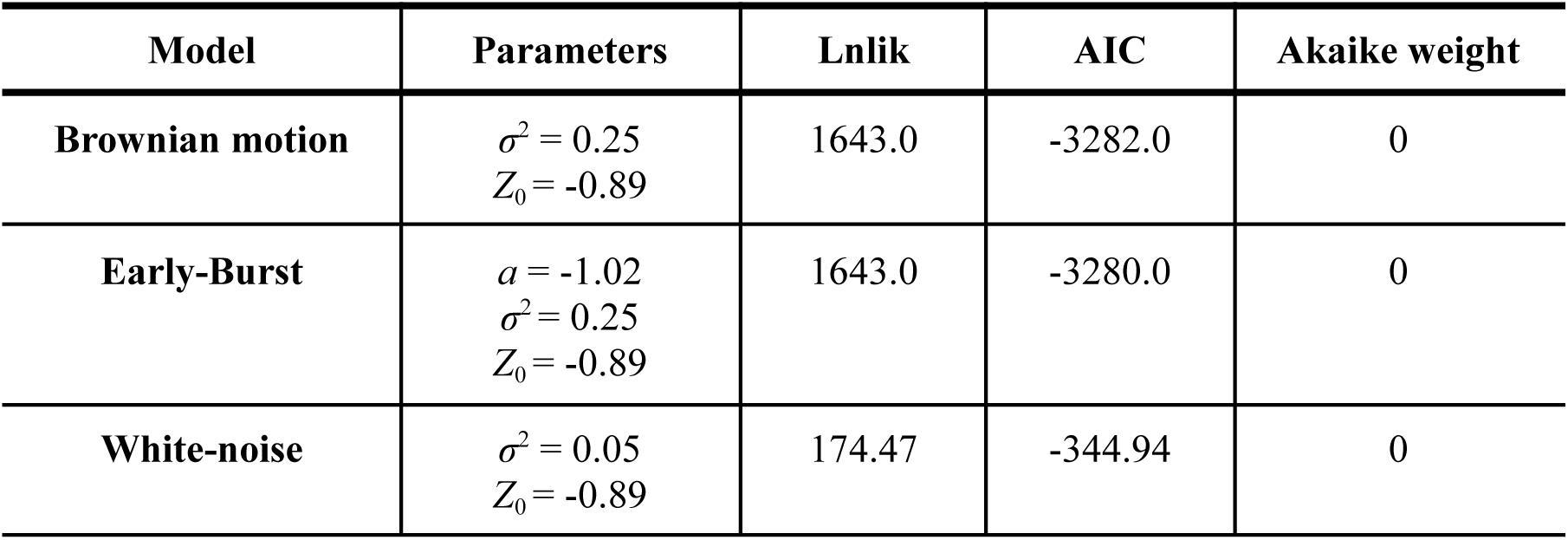
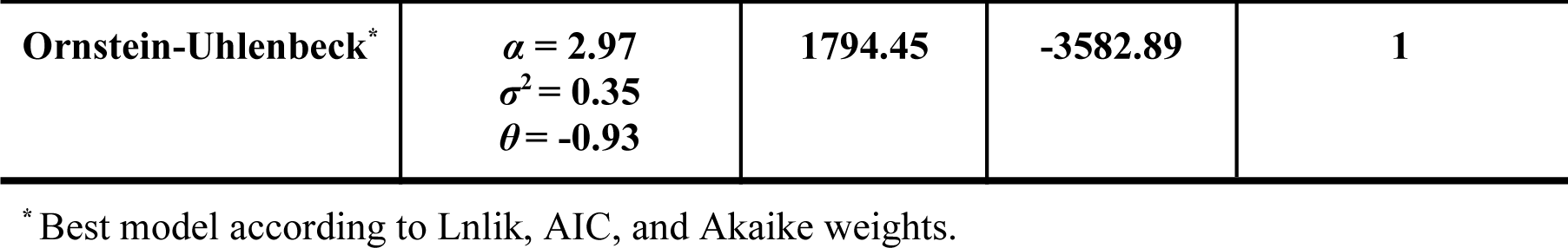
Summary of evolutionary models for cell size on a rescaled phylogenetic tree. OU model shows the highest likelihood and the smallest AIC. LnLik: log-likelihood, AIC: Akaike Information Criterion, *σ*^2^: rate of evolution of trait, *Z*_0_: trait value assigned at the root of the phylogenetic tree, *a:* pattern of rate decline, *α*: attractor strength, *θ:* optimum value.

| Model | Parameters | Lnlik | AIC | Akaike weight |
| --- | --- | --- | --- | --- |
| **Brownian motion** | $\sigma^2 = 0.25$<br>$Z_0 = -0.89$ | 1643.0 | -3282.0 | 0 |
| **Early-Burst** | $a = -1.02$<br>$\sigma^2 = 0.25$<br>$Z_0 = -0.89$ | 1643.0 | -3280.0 | 0 |
| **White-noise** | $\sigma^2 = 0.05$<br>$Z_0 = -0.89$ | 174.47 | -344.94 | 0 |
| **Ornstein-Uhlenbeck\*** | $\alpha = 2.97$<br>$\sigma^2 = 0.35$<br>$\theta = -0.93$ | **1794.45** | **-3582.89** | **1** |

\* Best model according to Lnlik, AIC, and Akaike weights.

### What forces may be behind a prokaryotic cell size optimum?

If one accepts that there is indeed a long-standing evolutionary pressure for prokaryotic cell size to have remained bound around such an optimum, it is only reasonable to ask what evolutionary pressures were at work, and if they may have left a recognizable imprint in the genomes of extant bacteria and archaea. To probe this possibility, and using available genomic metadata linked to 1361 species in our dataset, we assessed patterns between genomic traits and cell size. With respect to overall genome properties (genome size, number of proteins, GC content, coding density, number of 16S rRNA copies, and codon usage bias) we found a significant, if poor, correlation between log *V/S* and several genome traits (or their logs, as needed), with correlation values (*r_s_*) ranging from ™0.2 to 0.3 as assessed by Spearman’s rank correlation (**Figure S5**), ordinary least squares regression, and phylogenetic generalized least squares regression (**Figure S6, Table S6, S7)** to correct for phylogenetic dependence between traits ^49,50^. The situation did not improve by considering *V* instead of *V/S* as a measure of size (**Figure S7, Table S8, S9**). These results agree with those of a recent study that found a poor association between cell size and traits like genome size or maximum growth rate ^51^. In a similar but alternative exercise we also probed if particular KEGG database functional categories of genes attained differential importance in terms of either absolute or relative allocations of genomic length as a function of cell size, given that genomic complexity is associated strongly to functional traits providing significant selective advantages ^52^. Again here (**Table S10**) we found many functional categories whose genomic complexity was a statistically significant function of cell size, but also extremely weakly so in all cases. This type of blind, high-throughput analysis did not reveal any smoking-gun factor but rather suggests that a combination of factors may be responsible for the size optimum among bacterial cells. Finally, we carried out a few hypotheses-driven genomic analyses based on apparent trends observed in our dataset or on theoretical predictions from the literature regarding allometric relationships among bacteria, which are presented in **Figures S8, S9** and are addressed in detail in the discussion section.

## Discussion

### A typical prokaryotic cell

As noted previously ^1–3^, bacteria and archaea in our dataset span 3-6 orders of magnitude in both width and length. And yet, the quintessential bacterium comes as a rod-shaped cell with a linear dimension (*V/S*) of 0.1- 0.2 *μm*, a volume (*V*) of 0.1-2.1 *μm*^3^, and a surface area (*SA*) of 1.3-10.4 *μm*^2^. The probability that a bacterial species chosen at random fits this definition is slightly above 2/3. While spherical cells are not uncommon, it is unclear if they bring about particular adaptive value or if they merely represent evolutionary cul-de-sacs, given that rods can evolve into cocci by gene loss, but the opposite transitions are apparently not possible through linear descent ^53^. Interestingly the average archaeon is somewhat larger than the typical bacterium (Mann-Whitney test: *U* = 324195.5, *P-value* < 0.001): a range of *V/S* between 0.09-0.26 *μm* (with *V* between 0.18 and 2.89 *μm*^3^ and *SA* of 1.75-12.2 *μm*^2^) would encompass slightly above 2/3 of archaea in the dataset. Both rods and cocci are roughly equally represented among Archaea. Our dataset thus provides precise quantitative backing to the notion that prokaryotes are small-sized ^1^, even when some are outside of the common ranges, providing a measure to the sheer intensity of this size preference.

### Usefulness and limitations of the dataset

The dataset presented here can be useful in seeking evidence for biological hypotheses related to cell size by providing a large number of entries linked directly with minable genomic information. For example, bacterial genome size is commonly thought to co-vary with cell size. De Long et al., found an exponential relationship between genome size and cell volume with exponents around 0.35 (i.e., approximately scaling with linear size) using *n* = 28 species ^8^. Doubling the sample size (*n* = 57) reduced the exponent to 0.18 ^26^. Our current dataset (*n* = 1361; and see **Table S8** and **Figure S7**) indicates that genome size is clearly not a strong allometric function of cell size (the exponent fit is 0.07 ± 0.008 and *R^2^* = 0.05). Thus, the amount of genetic information required to run a bacterium seems largely independent of its size. We provide a few other examples of similar dataset uses in the discussion that follows below. And yet some caveats are also warranted. First, we do not address organismal size but cell size. The two are not always equivalent because multicellular bacteria do exist, even though they are not the norm. This complexity is not yet encompassed in our dataset or analyses, and it may have blurred some correlations or limited some conclusions. Second, our dataset is derived largely from cultured species, which may have introduced biases in phylogenetic coverage, as most cultured species tend to belong to a few phyla. However, it is unclear how large a bias this brings, because highly represented phyla like Proteobacteria are also often the most common and diverse in the natural environment. While we attempted to mitigate this bias by gathering data from multiple sources, some taxa will still be underrepresented, and most unculturable species are missing. However, sizes of uncultured species are well represented in our dataset as most cells from rare phyla and candidate divisions tend to have a spherical-equivalent diameter between 0.4 and 1.2 *µm* ^54^, corresponding to a log(*V/S*) range between ™1.2 and ™0.7. Additionally, if indeed standard microbiological cultivation techniques would be size-selective, this would add a systematic bias. While we could not find empirical demonstration for such an effect, one could argue that the fast-growing bacteria typically selected during enrichment and cultivation could be larger than the average, in keeping with some analyses ^23^. We discuss how this is likely inconsequential in the next section. Third, prokaryotic species do not exist in isolation but rather as part of communities with complex ecological interactions, which may impact cell size ^55,56^ in a manner not reflected in our dataset.

### A stability landscape for cell size

In accordance with previous work ^30,31^, our results demonstrate that cell size can be modeled as a diffusive process (**Table 3**). Building on this, we propose a stability landscape framework to understand the phenomenon. A simple approximation to such a stability function can be obtained as the difference between the empirical and log-normal distributions from Figure 2A (see Methods), which is presented in Figure 5. Within it, valleys represent basins of attraction, regions towards which the system gravitates. The depth of these basins represents barriers to overcome to change its state. The optimum cell size anchors the main basin of attraction. Species falling within this basin will tend to return to that optimum, regardless of phylogenetic history. Because the basin slopes are asymmetric, steeper towards smaller cell sizes than towards larger sizes, fewer barriers must exist for species around the optimum to become larger than smaller. Species outside two standard deviations from the optimum reach the basin’s edge, and fall within the influence of one of two secondary, shallower basins of attraction, one on either side. (though the depth of the left secondary basin may have been enhanced by the artifactual peak described above). The significance of these secondary basins, and the potential existence of secondary optima, is hard to assess precisely at this stage because of the low number of entries in these regions. As more genomic data becomes available, higher-resolution phylogenies could facilitate the exploration of multi-regime OU models to test the hypothesis of multiple optima.

**Figure 5.**
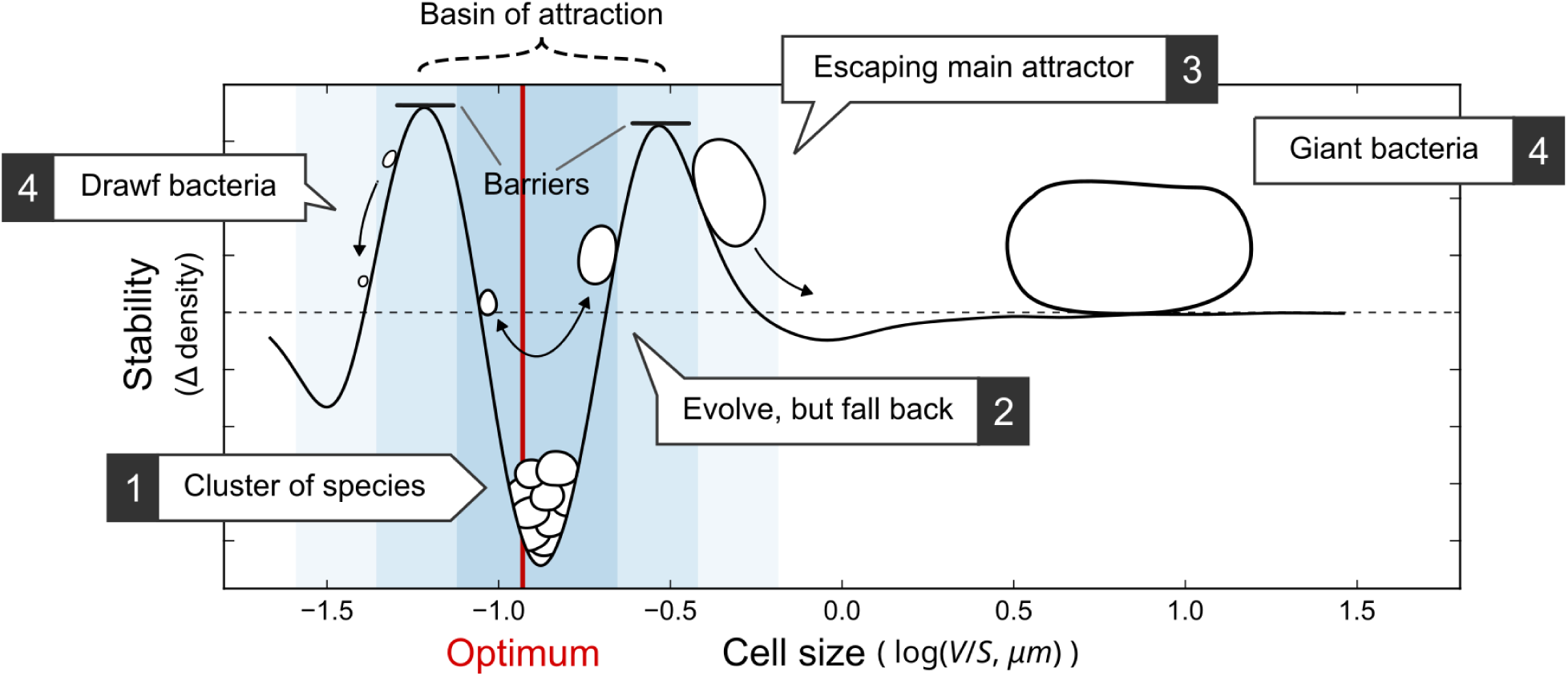
Stability landscape of cell size for prokaryotic species. There is a basin of attraction around the optimum cell size and most species fall within it. Very few species managed to escape the basin and acquire extremely large or small sizes. One, two, and three standard deviations from the mean are shown as a blue background gradient. Red line indicates the optimum cell size found by the OU model while dashed horizontal line indicates a Δ density value of zero.

### The necessity of an evolutionarily optimal size range

Our findings can only be interpreted in evolutionary terms by postulating that there exists a particular cell size conferring optimal fitness outcomes for most extant clades. However, we also saw that this optimum is not absolute, as exceptions do exist for at least two phyla (Cyanobacteria, which have a larger cell average cell size, and Tenericutes, which are significantly smaller) and for a few individual species that conform the groups of giant or dwarf bacteria. The immediate questions arising are: i) what factors determine this optimum, and ii) why precisely in this range. Potentially relevant factors can be either inherently cell biological, or a consequence of the physical environment, or even perhaps stem from complex microbial interactions, since they all demonstrably influence outcomes in which cell size can play a role. Based on both theory and empirical evidence, cell size is thought to play a major role in least constraining various organismal properties ^2,57–59^. While a comprehensive and appropriately quantitative evaluation of such factors is beyond our intent here, a discussion of the relevant literature in the light of our contributions is still warranted, particularly in attempting to explain the unimodal, close to log-normal size distribution of bacterial cell sizes.

### What evolutionary forces could keep bacterial cells from shrinking?

Because all life has minimal compositional requirements for viability and many cell components are not scalable, there must exist a minimal size for a cell ^22,28,60^. There will be a minimal number of genes required for basic cellular functions, for example, which will dictate a certain minimal DNA mass, and hence a certain minimal volume occupied by DNA ^60,61^. By contrast, bacterial membranes have largely invariant thickness and thus occupy an ever larger percentage of a cell’s total volume the smaller a cell is, constraining the relative space left for every other component,^62^. Just the consideration of membranes and DNA leads immediately to the existence of this limit. Theoretical derivations based on (more sophisticated) compositional and bioenergetic constraints suggest an absolute minimal cell volume range between 0.4 and 1 x 10^−2^ *µm*^3^^28^, a range only outdone by four bacteria in our dataset, but 2% of entries have in fact cell volumes within Kempes et al.’s bracket of minimal size. Dwarf bacteria have probably exhausted all shrinking possibilities. Approaching this lower limit, cells will experience an increasing loss of biological efficiency, as trade-offs are made in space allocation to various components. Additional potential drawbacks of becoming too small may include the fact that cellular quotas for particular components, like C content ^63^, DNA content ^26^ and size of the nucleoid ^27^ and cell membrane allotment ^24^ scale with cell volume with exponents less than one: the smaller a cell, the larger the relative investment needed per unit biomass. Because of the relevance of DNA and membrane phospholipids to the cellular phosphorus budget, cell phosphorus quotas are also likely higher in small cells. Because the cost of microbial motility scales negatively with size ^64^, exploring the environment should be costlier for smaller microbes relative to their size. And yet, a simple analysis of our dataset reveals a more nuanced picture: the fractional occurrence and genetic complexity of flagellar motility and chemotaxis genes, a proxy for its ecological/evolutionary relevance ^52^ among 1361 minable genomes, shows a maximum around (*V/S*) 0.13 *µm* (**Figure S8**).

### Could S/V ratios prevent bacteria from evolving cells much larger than the norm?

Potential benefits of a reduced cell size include, perhaps most famously, an increased efficiency for cell-surface dependent metabolic processes (such as respiration or nutrient uptake). Consistently, ATP synthase complex density in the bacterial membrane increases with cell size ^24^. And yet, modeling ^25^ suggests that a “respiratory deficit” occurs first when cocci exceed 1 *µm*^3^ or long rods grow beyond 10 *µm*^3^. If one considers doubling times of 10 hours, more typical in the natural environment, the threshold reaches 500 *µm*^3^. Combining all factors, it is possible to evade a respiratory limitation up to cell volumes of 10^5^ *µm*^3^^25^, which includes all but the four largest bacteria in our survey. If the *S/V* constraint were evolutionarily relevant, one would expect that fast-growing species would come in smaller sizes. Do they? In our hands, an analysis of all bacterial entries in the growth rate compilations of Lynch et al. ^23^, which includes those in previous surveys ^8,22^, does yield a negative allometric trend of growth rate with cell mass, in line with microorganisms as a whole ^23^, but this is very poorly predictive (*R^2^* = 0.09) and relatively insensitive to size (exponent fit = ™0.18 ± 0.05). Consistent with this, there seems to be no trace of correlation between cell size and genomic codon usage bias (CUB) in our dataset (**Table S6** and **Figure S6**). CUB is correlated with maximal growth rates in bacteria ^65^ and is used to estimate growth rate estimation from metagenomic data in microbial ecology ^66^. Additionally, because rods have a more favorable shape than cocci to deal with surface limitation, one would expect that their average size could be larger, but we found just the opposite (Figure 1). Finally, an examination of size distributions in *Lactobacilli* in our dataset is useful. With very few exceptions ^67^, *Lactobacilli* gain energy exclusively through substrate-level phosphorylation, have no membrane electron transport carriers, and should be immune to *S/V* ratio limitations to energy generation, and hence to any subsidiary limitations to cell size. But we find no significant difference between average log cell size of obligately fermentative *Lactobacilli* (*n* = 126, log *S/V* = ™0.05 ± 0.50) and that from the rest of the Firmicutes in our survey (*n* = 666; log *S/V* = 0.05 ± 0.51), a 2-tailed, equal variance T-test yielding *P-value* = 0.93. In sum, any relationship between cell size and growth rate that might exist is weak and forgiving enough to allow for high variability and low predictability. Similarly, there is no empirical evidence relating cell size to increased nutrient uptake efficiency among bacteria, and experimental rates of nutrient uptake are not sensitive to cell size in phytoplankton (including bacteria and much larger eukaryotic species) ^68^. While in specific situations *S/V* ratio effects could have left an imprint in the evolution of specialized bacterial types (extremely fast growers, extreme oligotrophs), it is not plausible that such effects would invariably and necessarily be a metabolic burden that would result in the progressive paucity of ever larger bacteria. Unusual structural traits such as internal vacuolization that are consistent with a limitation in substrate supply, are reported only in giant bacteria ^9,69^.

### What other forces could moderate cell size?

Other non-metabolic potential benefits to be derived from a smaller cell may include an ability to escape bacterivorous flagellates ^14^, viral attack ^70^, or enable propagation through porous media, as exemplified by the development of tiny cyanobacterial baeocytes in endoliths ^71^. This does not quite seem to offer a universal driver either. One could also consider purely physical forces that vary with linear dimension. While for phototrophs an increase in size taxes the efficiency of light absorption through self-shading ^72^, it is precisely the purely phototrophic phylum (Cyanobacteria) that has the largest average size of all (Figure 2B). Finally, and we think importantly, mass transport is another scale-sensitive, relevant process. At small scales, viscous forces dominate over inertial forces, the environment turning essentially stagnant. Without mass transport by turbulent (or organized) flow, diffusion becomes its main vehicle. Average diffusion times scale with L^2^ ^74^. A small molecule such as sucrose diffuses across a microbial coccus 1 *µm* in diameter in about a millisecond but requires 15 minutes to do so through a 1 *mm* cell. A ribosome will make it across one micrometer in less than a second but will need half an hour to cross a 100 *µm* cell. These are likely underestimates since molecular mobility in the cytoplasm is subdiffusive ^75,76^, particularly for large molecules ^77^. There are also consequences for regulatory processes: phosphorylation decay times being invariant, signaling will be localized rather than global as cells grow beyond a few *µm* in diameter ^78^. In the absence of the ATP-dependent cytoplasmic “active” transport or cytoplasmic streaming of eukaryotes, diffusion limitation to cellular physiology, regulation and metabolism are likely an evolutionary deterrent to cell size increase in bacteria and archaea ^76,77^, one whose effects should be universal. If diffusive limitations were at play and noting that spheres constitute the most effective shape for diffusive content homogenization by minimizing average paths ^74^, one could predict that the average size of cocci could be larger than that of other bacterial cell shapes, as we indeed found (Figure 1). In our view, a direct assessment of the role of diffusion in preventing prokaryotes from becoming larger (while allowing eukaryotes to do so) would be desirable. To curb one’s enthusiasm, the existing negative allometric relationship between cell size and growth rate discussed in the previous paragraph, while principally consistent with the notion, is much weaker than could be expected from a major driver.

### Autotrophs vs. heterotrophs

One of the salient patterns we could discern from our dataset is the fact that autotrophs, which make up only a small proportion of bacterial and archaeal species overall, are well represented among large species. Two of the four “giant” bacteria in our dataset, and 71 the 100 largest bacteria in our dataset are autotrophs. This clearly exceeds the representation of autotrophs among cultivated prokaryotes. The enrichment of autotrophs among large-celled species can also be shown by data mining for the incidence of Rubisco in our dataset (**Figure S9**). This finding can hardly be stochastic. What distinguishes autotrophs from heterotrophs is a reliance on CO_2_ as a source of cellular carbon for growth. Typically, autotrophs have slower growth rates since all building blocks must be synthesized de novo. Broad differences in generation time could explain the pattern, in that inherently longer doubling times could counter limitations in respiration, nutrient uptake or intracellular diffusion of molecules and signals. If the limitation to expand cell size were based on diffusion of cellular components as discussed in the previous paragraph, then an autotroph’s inherently longer generation times could accommodate larger cell sizes than faster-growing heterotrophs.

### A multiplicity of drivers

The preceding discussion of potential drivers leads us to favor the concept of cell size as an emergent trait shaped by multiple interacting drivers. In his perspective, Koch ^79,80^ proposed two mechanisms that may result in log-normal distributions of size through multiplicative processes. The first mechanism suggests that size approximates log-normality as the product of two correlated linear dimensions, such as length and width. However, for prokaryotes, any measure of size (length, width, volume, or surface area, *V/S*, *S/V*) yields a distribution with the same properties (**Figure S10** and **Table S11**), thus challenging this idea. The alternative mechanism suggests that fluctuations in the trait arise through multiplicative interactions of multiple factors. This concept has been integrated into a macro-evolutionary model that successfully replicates the empirical body size distribution of mammals as a cladogenetic (speciation and extinction) multiplicative diffusion process over evolutionary time ^30^. Further emphasizing this complexity, our preliminary regression analysis indicated that at least 25 of 269 features, including genome properties and KEGG metabolic pathways, were needed to account for close to half of the variance (*R^2^* = 0.44; **Figure S11**). Thus, the most parsimonious explanation in our view is that cell size is an emergent trait with remarkable plasticity yet highly conserved across time, where no one driver is sufficient to explain the diversity of cell sizes observed in the prokaryotic world, but drivers interact to define size optima.

## Acknowledgments

We are grateful to Dr. Michael Lynch and Dr. Stephanie Forrest for co-organizing a symposion on cell size with F.G.-P., where we discussed our work with colleagues and received valuable feedback. We are indebted to Ángel Robles-Fernández for valuable discussions on initial analyses. This research was partly supported by an Arizona State University start-up grant to Q.Z. and a grant (DEB 2129537) from the National Science Foundation to F.G.-P..

## Author contributions

F.G.-P. and Q.Z. conceived the study. H.S.-M. led data collection and curation, data analysis, result interpretation and manuscript writing. A.C. and L.G.-S. contributed to data collection and curation. F.G.-P. led advisory of biological discussion and contributed to data analysis. Q.Z. led advisory of data analysis and contributed to biological discussion. All co-authors contributed to the writing and discussion of the manuscript.

## Declaration of interests

The authors declare no competing interests.

## Methods

### Data collection and curation

We collected the data of cell length, width, and shape of bacterial and archaeal species from four sources: Bacterial Diversity (BacDive) Metadabase on November 2021 ^81^, Pasteur Culture Collection on December 2021 ^82^, and the Bergey’s Manual of Systematics of Archaea and Bacteria Second Edition on July 2022 ^16^. This multi-source approach was adopted to maximize the diversity of species included in the dataset. We then curated the values of cell size from each source. This entailed fixing typos in the reported values and consolidating units to *μm*. For species reported as sphere-like with diameter values for one dimension, we set cell width equal to length and equal to reported diameter. Species with non-spherical cells whose reported size was only in one dimension were discarded. When values were reported as ranges, we calculated the geometric mean of reported minima and maxima. For example, 0.5–2.0 *μm* was entered as 1.0 *μm*, whereas ≥ 0.5 *μm* became 0.5 *μm*. This all resulted in single values assigned to single species. Cell volume and surface area for each species were then calculated using the formulae for a capsule (i.e., a rod):

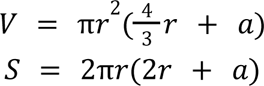

where 2*r* = width and *a* = length − width. For spherical cells *a* becomes zero, and the formulae reduce to those for a sphere. Hence, these formulae estimate the volume and surface area for rod and spherical shapes, but only approximate them for other shapes. As a standard, shape-independent one-dimensional measure of cell size we calculated and used the ratio of cell volume to surface, *V/S*. A mapping between length, width and *V/S* for rod and spherical cells is shown in **Figure S12**.

### Phylogenetic analysis

We assigned species in our dataset to a unique taxonomic identifier (TaxID) at the species level using a local version of the NCBI taxonomy database updated in November 2021 ^83^. Conflicts and ambiguities were manually resolved using the NCBI taxonomy browser. This enabled us to classify species into the six standard taxonomic ranks (species, genus, family, order, class, and phylum). For records that shared the same taxonomic identifier, we calculated the geometric mean of cell length and cell width area to obtain a single number. The resources from the Web of Life (WoL) project ^41^ were used for phylogenetic analysis. Specifically, we used the reference phylogeny annotated with tax2tree ^84^, which allowed us to map the species taxonomic identifiers from our dataset to nodes of the WoL reference phylogeny. The mapping step consisted of assigning the species to the lowest ancestor node that contains the species. Once the mapping was completed, we attached the species as a tip to the mapped node with a branch length equal to the median depth of all descendants of the node. Finally, we pruned the phylogenetic tree such that it only contains the species from our dataset that were inserted, thus obtaining the phylogenetic tree used for the presented analyses.

### Genome properties, codon usage bias, and optimal growth temperature

The WoL project contains metadata associated with genome properties, including genome size, number of proteins, coding density, GC content, and number of 16S rRNA copies. For species that were mapped at the tips of the WoL reference phylogeny, the genome properties were inherited from the WoL metadata. The codon usage bias was estimated as the Effective Number of Codons given the G+C compositions (ENC’), following the procedure described in ^65^. For this, we first identified the set of highly expressed genes (HEGs) for the 10,575 genomes from the WoL using a local version of the Dynamic Codon Biaser tool ^85^. We then calculated the value of ENC’ for the concatenation of all coding sequences (ENC’_all_) and the concatenation of HEGs (ENC’_HEGs_) with the function ENCprime from the R package CoRdon v1.12.0 ^86^, using the bacterial, archaeal, and plant plastid code. Finally, the ΔENC’ was calculated as:

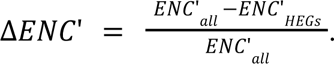

We retrieved the average optimum growth temperature from the TEMPURA database ^87^ on September 2022 by mapping the TaxID of each species in our dataset to the identifier in TEMPURA.

### Correlation between cell size and phylogenetic distances

To assess whether differences in cell size among species correlates with evolutionary distance, we calculated 1) the patristic distance (i.e. tip-to-tip distance) of the phylogenetic tree and 2) the difference in log(*V/S*) for every pair of species, which resulted in C(5,380; 2) = 14,469,510 pairs. Because some pairs included species with extremely different sizes (e.g. *M. genitalium* and *T. namibiensis*), we normalized the difference in size as

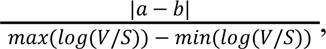

where a and b are the sizes of the two species, respectively. We refer to this normalization as the Size Difference Index (SDI) in log(*V/S*). The normalization step produces values bounded between 0 and 1, where smaller values indicate similar sizes and vice-versa. The relationship between size and evolutionary distance was assessed with polynomial fitting, exponential fitting, linear regression, and piecewise linear regression (see below for details). Moreover, we calculated the goodness of fit for each model with the coefficient of variation (*R^2^*). Only the model with the best fit is shown in Results. Given the non-homogeneous distribution of species pairs across evolutionary distances, we binned them into distance ranges in order to increase the signal-to-noise ratio and capture the emergent statistical pattern. Specifically, we chose a bin size of 0.2 and randomly sampled 1000 pairs for each bin. Values of bin size and number of species sampled per bin do not affect the overall pattern. To determine the taxonomic levels corresponding to different ranges of patristic distances, we employed the following approach. For every species pair, we asked at what patristic distance each species is classified into different taxonomic groups. For instance, let us consider a pair of species belonging to different families. We start asking whether they differ at the phylum level. Finding no distinction, we then proceeded to the class level and repeated the inquiry until reaching the family level. As the species pair differed at the family level, we stored their patristic distance into a vector, which was later used to estimate the distribution of distances for all species differing at the family level. The application of this iterative approach enabled us to determine the corresponding taxonomic levels associated with specific ranges of patristic distances.

### Phylogenetic signal and evolutionary models

We quantified the phylogenetic signal for cell size, genome properties, codon usage bias, and average optimal growth temperature using the function fitContinuous from the R package geiger v2.0.7 ^88^ for Pagel’s *λ* and the function phylosig from the R package phytools v1.0.1 ^89^ for Blomberg’s *K*. We used a likelihood ratio test to assess the statistical significance of rejecting the null hypothesis of no phylogenetic signal (i.e. *λ* = 0) for Pagel’s *λ*. For Blomberg’s *K*, we assessed the statistical significance with a permutation test of 1000 simulations. To quantify the phylogenetic signal across taxonomic groups, we pruned the phylogenetic tree such that it only contained the species in the group of interest and estimated Pagel’s *λ* as described above. After quantifying the phylogenetic signal, we fitted and compared four models of trait evolution, including Brownian motion ^42^, Early-Burst ^45^, White-noise, and Ornstein–Uhlenbeck ^46^ for *V/S*. We used the fitContinuous function with default parameters from the geiger package, and we selected the model that best fits our data with the aic.w function that estimates the relative contribution of each model using Akaike weights ^47^. We tested the four models of evolution in phylogenetic trees with and without rescaling their height to 1 for interpretation purposes (see Results).

### Statistical analyses, linear regressions, phylogenetic regressions, and polynomial fitting

We log-transformed *V/S* to reduce the right-skewness caused by larger cells and used this metric for all subsequent analyses. Species were binned into discrete ranges of log(*V/S*) with the histogram function from the Python package NumPy v1.20.3 ^90^. The optimal bin size was chosen automatically (as the larger of the Sturges and Freedman Diaconis estimators). This enabled the estimation of cell sizes distributions. We also performed kernel density estimation (KDE) to smooth the distribution across phyla using scikit-learn v0.23.2 ^91^. The optimal bandwidth for KDE was chosen by hyperparameter tuning through 5-fold cross-validation. To evaluate how well the distributions approximate a log-normal distribution, we conducted a goodness-of-fit analysis. For this, we performed a Kolmogorov-Smirnov test between the values of log(*V/S*) and a log-normal distribution with log(*μ*) and log(*σ*) estimated from the data. Values of skewness and kurtosis were also estimated from the distributions. Likewise, we generated quantile-quantile plots to better evaluate the deviations from log-normality. Because *V/S* does not follow a log-normal distribution for Proteobacteria, Firmicutes, and Tenericutes, we used the non-parametric Kruskal-Wallis followed by Mann-Whitney *U* tests to compare the values of cell size across phyla. In addition, we evaluated the homogeneity of variances across taxonomic groups with the Levene’s test. We then corrected the reported *P-values* of all statistical tests using the Benjamin-Hochberg method. All these statistical analyses were carried out using functions from the Python package SciPy v1.6.2 ^92^. We used Spearman’s rank correlation coefficient to assess the correlation between log(*V/S*) or log(*V*) and genome properties. We also estimated the predictive power of cell size on genome properties with linear regressions using the OLS function from the Python package statsmodels v0.12.0 ^93^. We controlled for phylogenetic non-independence in the residuals using a phylogenetic generalized least squares method implemented in the pgls function from the R package caper v1.0.1 ^94^.

### Approximation of the stability landscape

Because we do not have a set of equations describing the dynamics of cell size, we approximated the derivation of the stability landscape. We used the empirical distribution from Figure 2 because it contains the actual dynamics of species-specific cell sizes sampled at the current time, including the bias towards larger sizes and tendency of species to cluster around the optimal size. The log-normal distribution fitted to the data can serve as a null model because it does not include any bias or optimal value. The stability landscape was then derived as the difference between the probability density functions of the two distributions (Δ density) as it highlights the differences between the observed and expected dynamics of a multiplicative null model. Thus, the steepness of the basins represents deviations from the null model with positive values indicating less species than expected and negative values indicating more species than expected.

### Relationship between cell size and genome function

Leveraging the genome annotations available in the WoL project, we assessed whether species contain genes necessary in photosynthesis, chemotaxis, and flagellar movement. We used annotations from the KEGG database, which annotates coding sequences into functional orthologs based on homology and group orthologs into pathways ^95^. For example, the KEGG photosynthesis pathway has 63 orthologs assigned. Therefore, to assess whether a bacterial species can perform photosynthesis, we first counted the number of different genes in the pathway. Because not all the genes involved in photosynthesis are conserved across species, we decided to use a threshold in the number of genes that a species contains to classify the species as either positive or negative for photosynthesis. This threshold introduces flexibility in the number of genes present and ensures a more accurate representation of the species’ function. For instance, a high threshold represents a scenario in which the number of core genes for the pathway is high, whereas a lower threshold is a scenario in which the pathway contains a low number of core genes. We used a threshold of 0.7 to test for photosynthesis, flagellar assembly, and chemotaxis.

A more granular approach to assess species functional capabilities is to use a single gene that is essential for proper function. Take for example, the rubisco enzyme, used by autotrophs to convert CO_2_ into glucose, which is represented in the KEGG database by two orthologs (K01601: large chain, K01602: small chain). Species containing these two orthologs can then be classified as photosynthetic. To further assess the relationship between cell size and species function, we binned cell sizes into discrete bins of size 0.2. We combined those bins that contained less than 10 species. For each bin, we then calculated the fraction of species classified as positive for the respective pathways. In an alternative exercise, for each species, we quantified the relevance of KEGG pathways for cell size using four different metrics ^52^: the different number of proteins, the total number of proteins, the total number of base pairs of all genes in a pathway, and the fraction of total number of base pairs of all genes divided by genome size. We then fitted OLS regressions and calculated the scaling exponent and coefficient of determination.

### Supervised prediction of cell size based on genomic information

We compiled 269 features, encompassing six genomic properties (genome size, GC content, number of proteins, number of 16S rRNA genes, codon usage bias, and coding density) and 263 KEGG metabolic pathways. Logarithm transformation was applied to genome size and number of proteins, and arcsine transformation to GC content and coding density. The fraction of total number of base pairs of all member genes divided by the genome size was used as a measure of ecological/evolutionary relevance of a pathway. Supervised regression was performed using scikit-learn v1.3.2 ^91^. The dataset consisting of 1361 samples was split into training and test sets, with a ratio of 7:3. A pipeline was constructed, consisting of standardization followed by fitting with a random forest regression model, using default parameters. The 269 features were iteratively reduced using recursive feature elimination with cross-validation (RFECV). Specifically, in each iteration, one feature that has the lowest importance score as calculated by the random forest regressor was removed from the feature set, and the pipeline was executed with 10 repeats of 5-fold cross-validation on the training set to assess model performance as indicated by the coefficient of determination (*R*^2^). A saturation curve of model performance vs. feature count was plotted and manually examined to determine where model performance plateaued. The feature set with the highest model performance was retained and used to train a final model, whose performance was evaluated against the test set.

## Resource availability

### Lead contact

Further information and requests for resources should be directed to and will be fulfilled by the lead contact, Qiyun Zhu.

### Materials availability

This study did not generate new unique reagents.

### Data and code availability

This paper analyzes existing, publicly available data. The data sources and the procedures for data collection and curation are detailed in the “Data collection and curation” section.

The original and curated data, the analytical workflows (including Python, R and Linux scripts, with appropriate documentation), and the results of analyses presented in this paper are publicly available at: https://github.com/HSecaira/ProkaCellSize.

Any additional information required to reanalyze the data reported in this paper is available from the lead contact upon request.

## Supplemental information

Supplemental figures S1-S11.

Supplemental tables S1-S11.

## Notes

https://github.com/HSecaira/ProkaCellSize

### Competing Interest Statement

The authors have declared no competing interest.

