## Supplemental figures and tables for "An evolutionary optimum amid moderate heritability in prokaryotic cell size": supplemental_data.docx

### **Supplemental information**

#### **Supplemental figures**

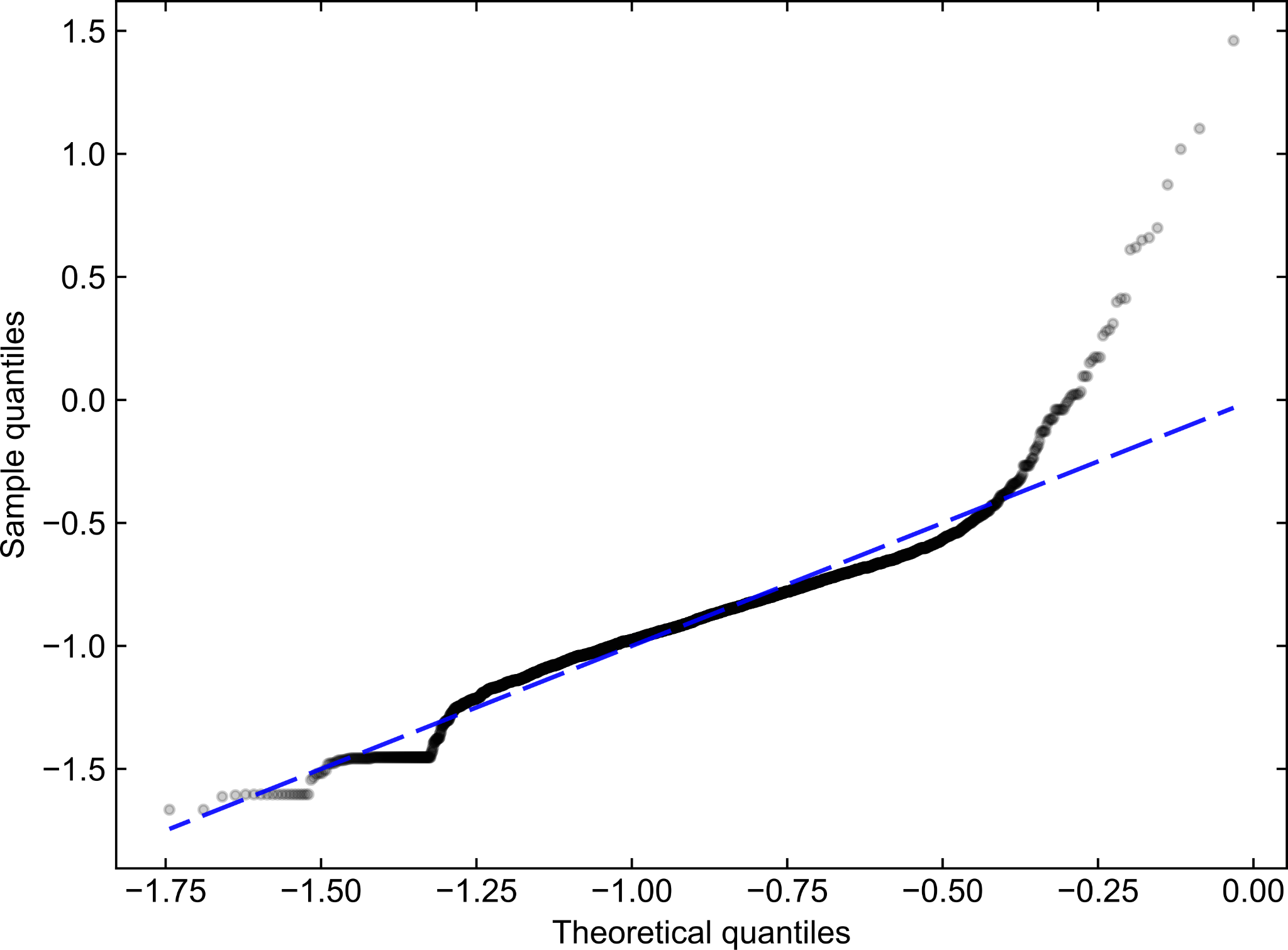

**Figure S1. Related to Figure 2A. Distribution of cell sizes does not follow a log-normal distribution.** Quantile-quantile (QQ) plot between the empirical distribution of volume-to-surface (*V/S*) ratios (black dots) and log-normal distribution (blue dash/dot line) fitted to the dataset. Extremely small and large ratios produce a high density of outliers (leptokurtosis), while large ratios alone produce right-skewness.

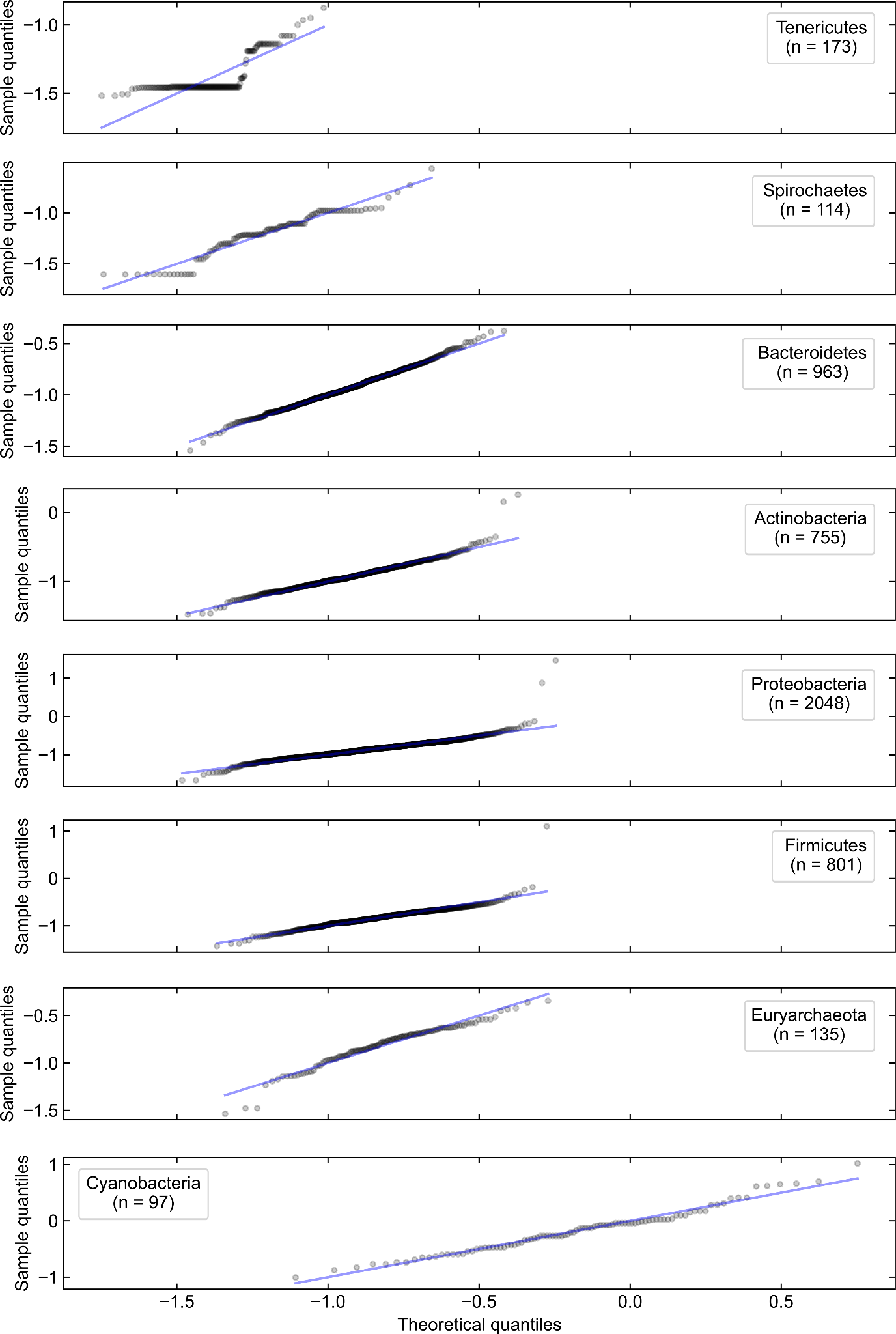

**Figure S2. Related to Figure 2B. Empirical distributions of cell size across phyla follow a log-normal distribution except for Proteobacteria, Firmicutes, and Tenericutes.** QQ plots between the empirical distributions of *V/S* (black dots) and log-normal distributions (blue dash/dot line) fitted to the data. Log-normal distributions were fit to the data using the mean and standard deviation estimated from each phyla. Tenericutes show a straight horizontal line, which is reflected as a peak in their cell size distribution.

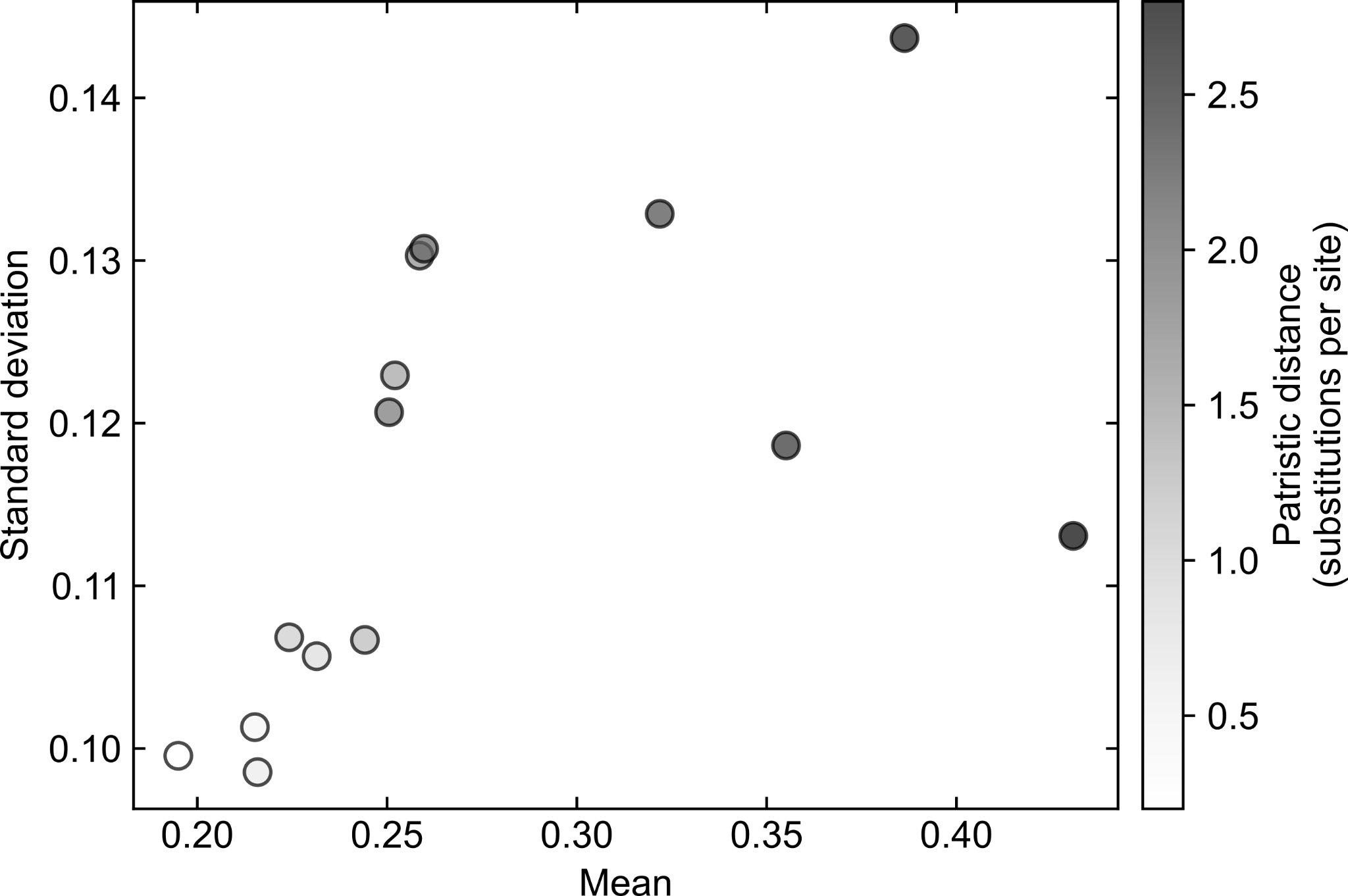

**Figure S3. Related to Figure 3.** Increase in mean and standard deviation across distance-binned size divergence index distributions.

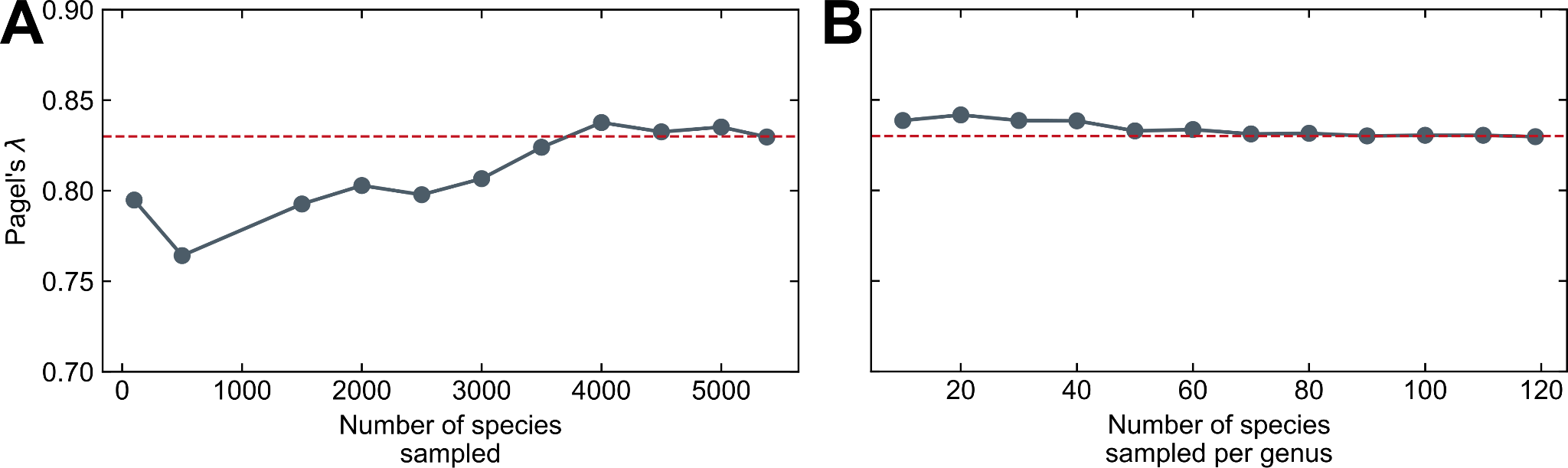

**Figure S4. Related to section “Heritability of prokaryotic cell size”.** Robustness in the estimation of phylogenetic signal for cell size (*V/S*) as a function of sampling efforts, in absolute (A) or relative (B) terms. Red dashed line represents the value estimated of Pagel’s *λ* = 0.83 for all species in our dataset.

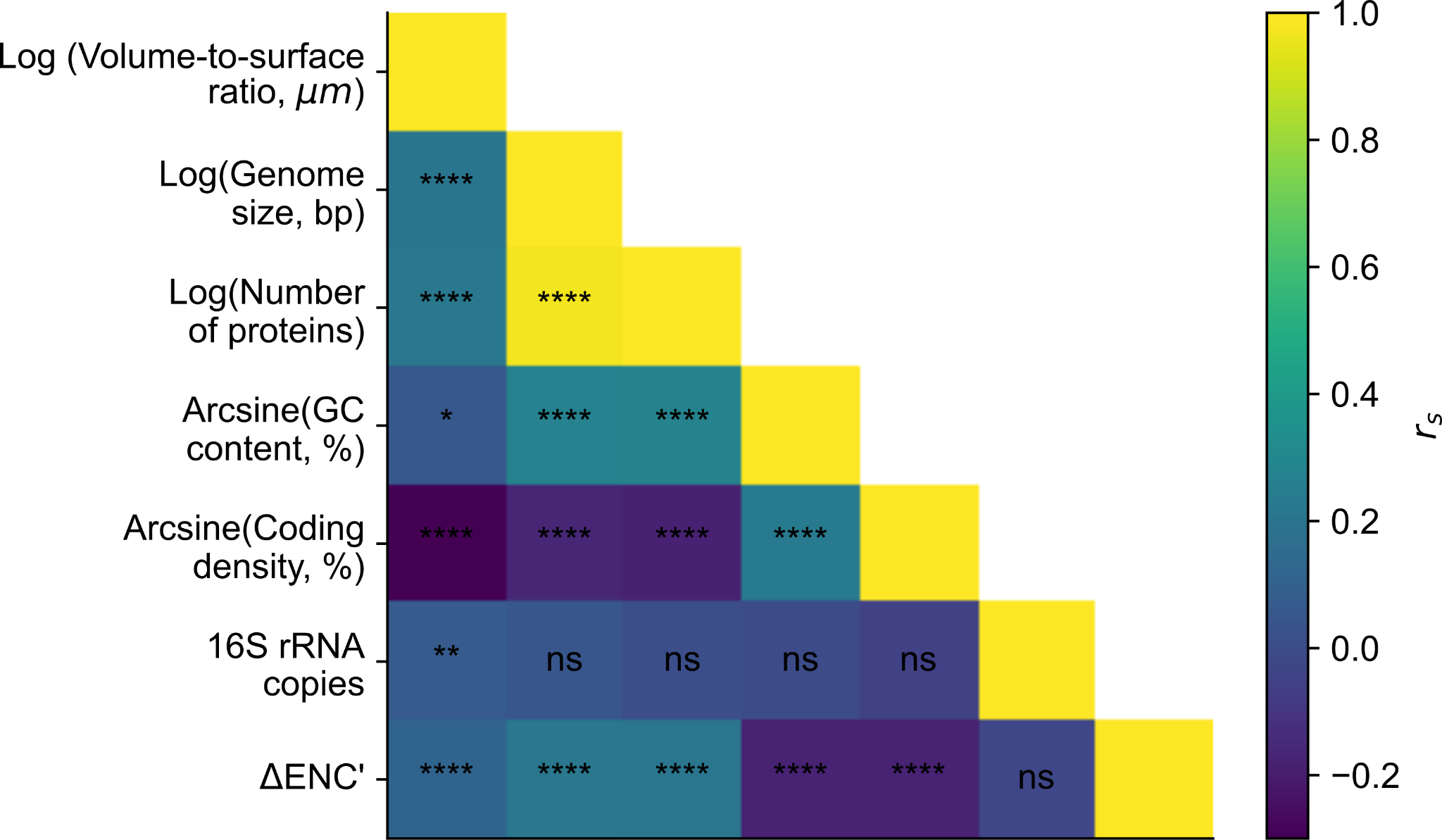

**Figure S5. Related to section “What forces may be behind a prokaryotic cell size optimum?”.** Poor correlation between genome traits and cell size. Spearman’s rank correlation coefficient (*r_s_*) between genome traits and volume-to-surface ratio. Significance levels: ns: *P*-value > 0.05, *: *P*-value ≤ 0.05, **: *P*-value ≤ 0.01, ***: *P*-value ≤ 0.001, ****: *P*-value ≤ 0.0001.

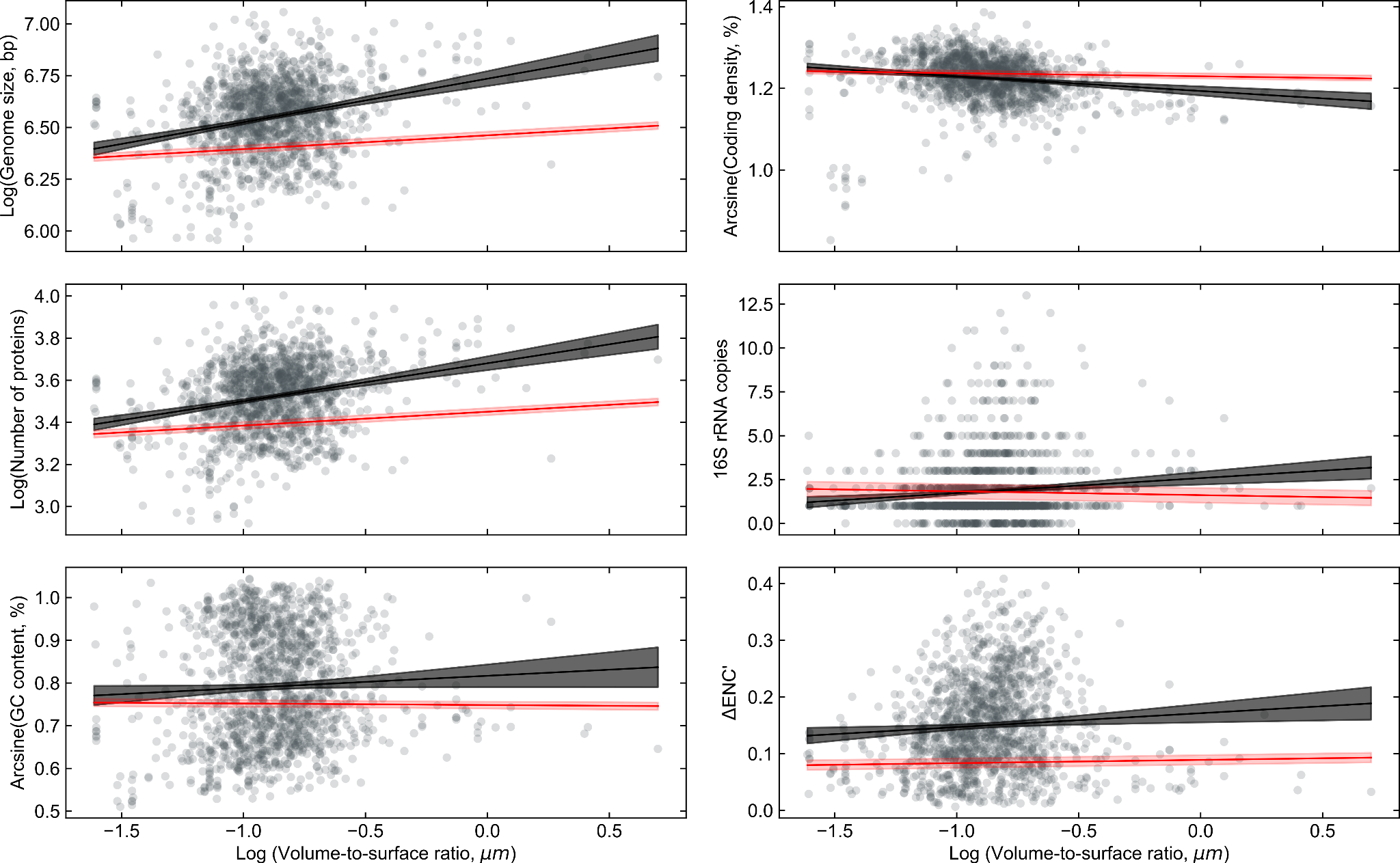

**Figure S6. Related to sections “What forces may be behind a prokaryotic cell size optimum?” and “Could S/V ratios prevent bacteria from evolving cells much larger than the norm?”.** Relationship between cell size (volume-to-surface ratio) and genome traits. Black line indicates the best linear fit according to OLS and shaded area is the 95% confidence interval. While red line is the best fit according to PGLS and shaded area is the 96% confidence interval. Gray points represent individual species.

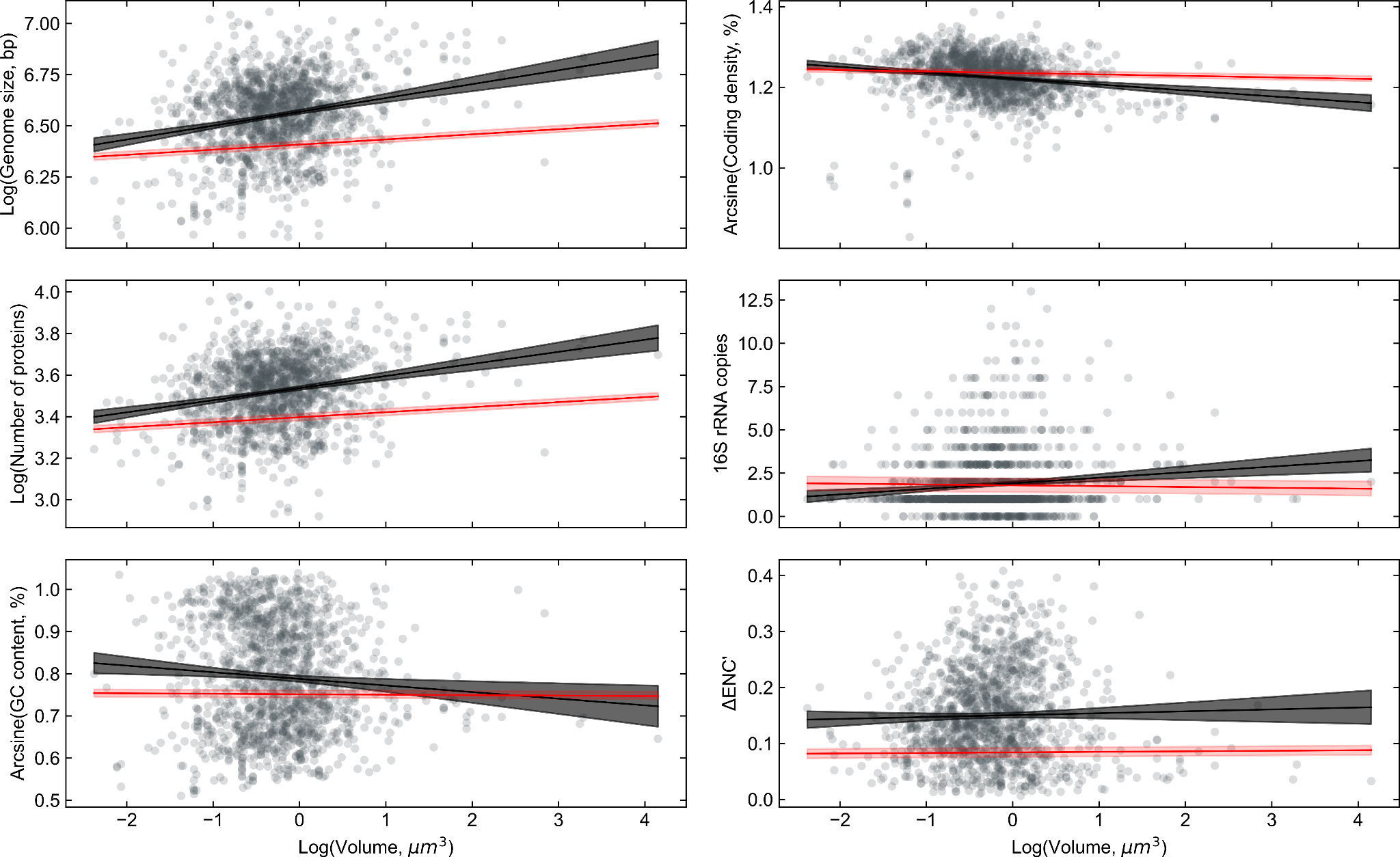

**Figure S7. Related to sections “What forces may be behind a prokaryotic cell size optimum?” and “Usefulness and limitations of the dataset”.** Relationship between cell size (volume) and genome traits. Black line indicates the best linear fit according to OLS and shaded area is the 95% confidence interval. While red line is the best fit according to PGLS and shaded area is the 96% confidence interval. Gray points represent individual species.

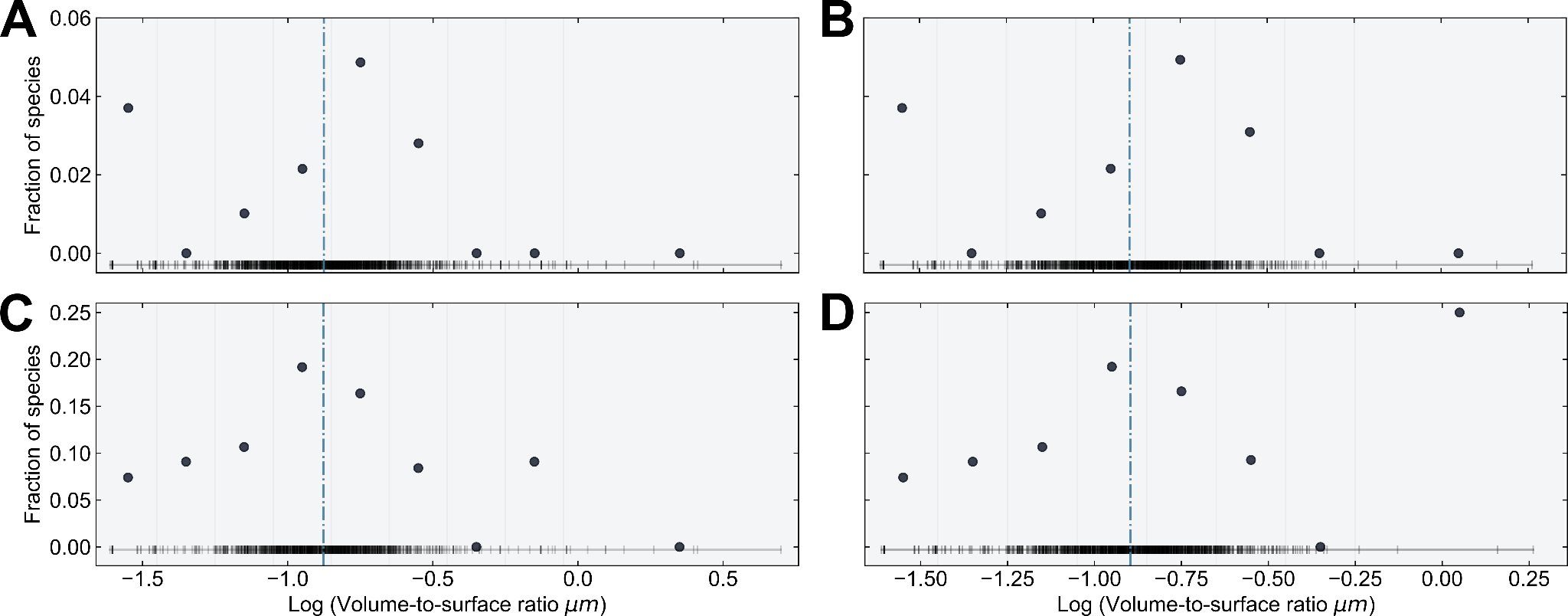

**Figure S8. Related to sections “What forces may be behind a prokaryotic cell size optimum?” and “What evolutionary forces could keep bacterial cells from shrinking?”.** Fraction of species containing at least 70% of genes involved in chemotaxis (A, B) and flagellar assembly (C, D) pathways. Left panels contain all species (*n* = 1361), while right panels contain all but Cyanobacteria species (*n* = 1,316). Black dots represent the fraction of species in bins (gray vertical lines), while short vertical black lines close to the abscissa indicate individual entries. Blue dashdot line indicates the mean of the *V/S* distribution.

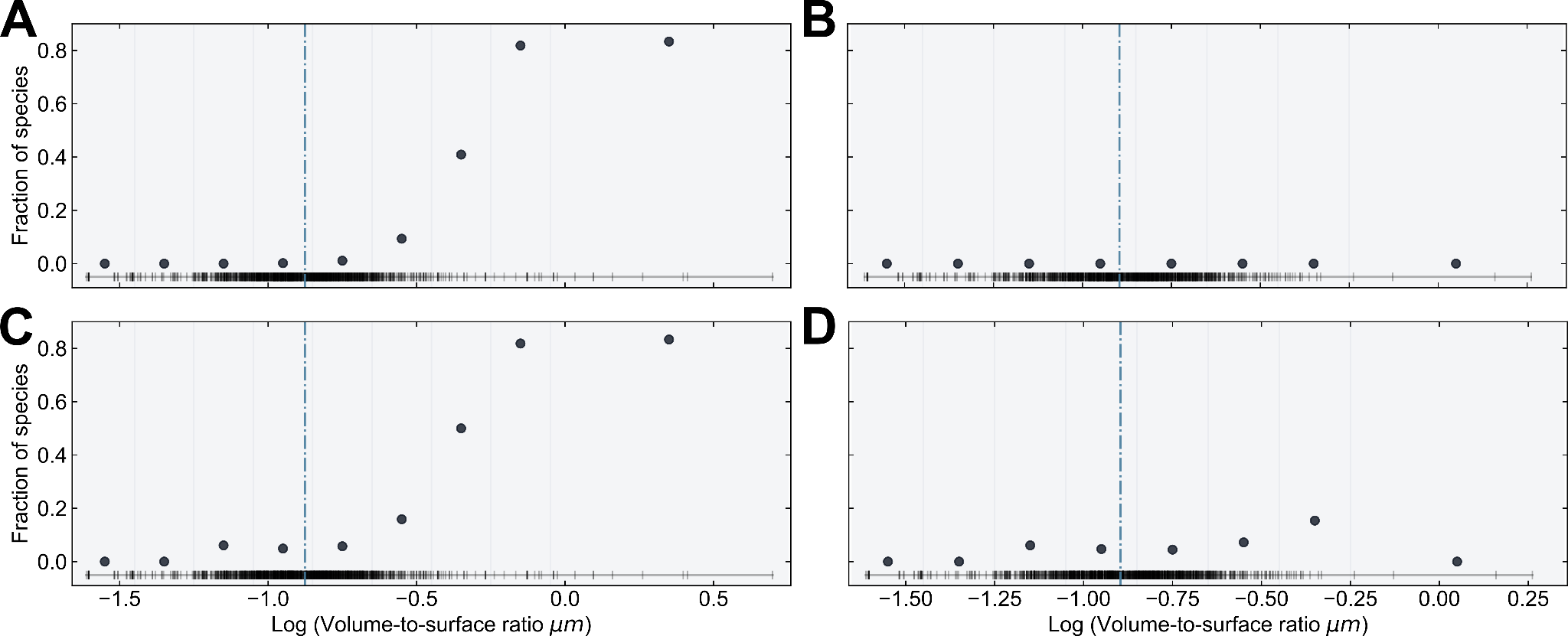

**Figure S9. Related to sections “What forces may be behind a prokaryotic cell size optimum?” and “Autotrophs vs. heterotrophs”.** Fraction of species containing at least 70% of genes involved in photosynthesis pathway (A, B) or the two subunits of Rubisco (C, D). Left panels contain all species (*n* = 1361), while right panels contain all but Cyanobacteria species (*n* = 1,316). Black dots represent the fraction of species in bins (gray vertical lines), while short vertical black lines close to the abscissa indicate individual entries. Blue dashdot line indicates the mean of the *V/S* distribution. Notice that if Cyanobacteria species are removed, only the analyses of Rubisco yields a fraction of species higher than zero, thus highlighting that our analyses are sensible to the level of granularity (i.e. single genes or set of genes in pathway).

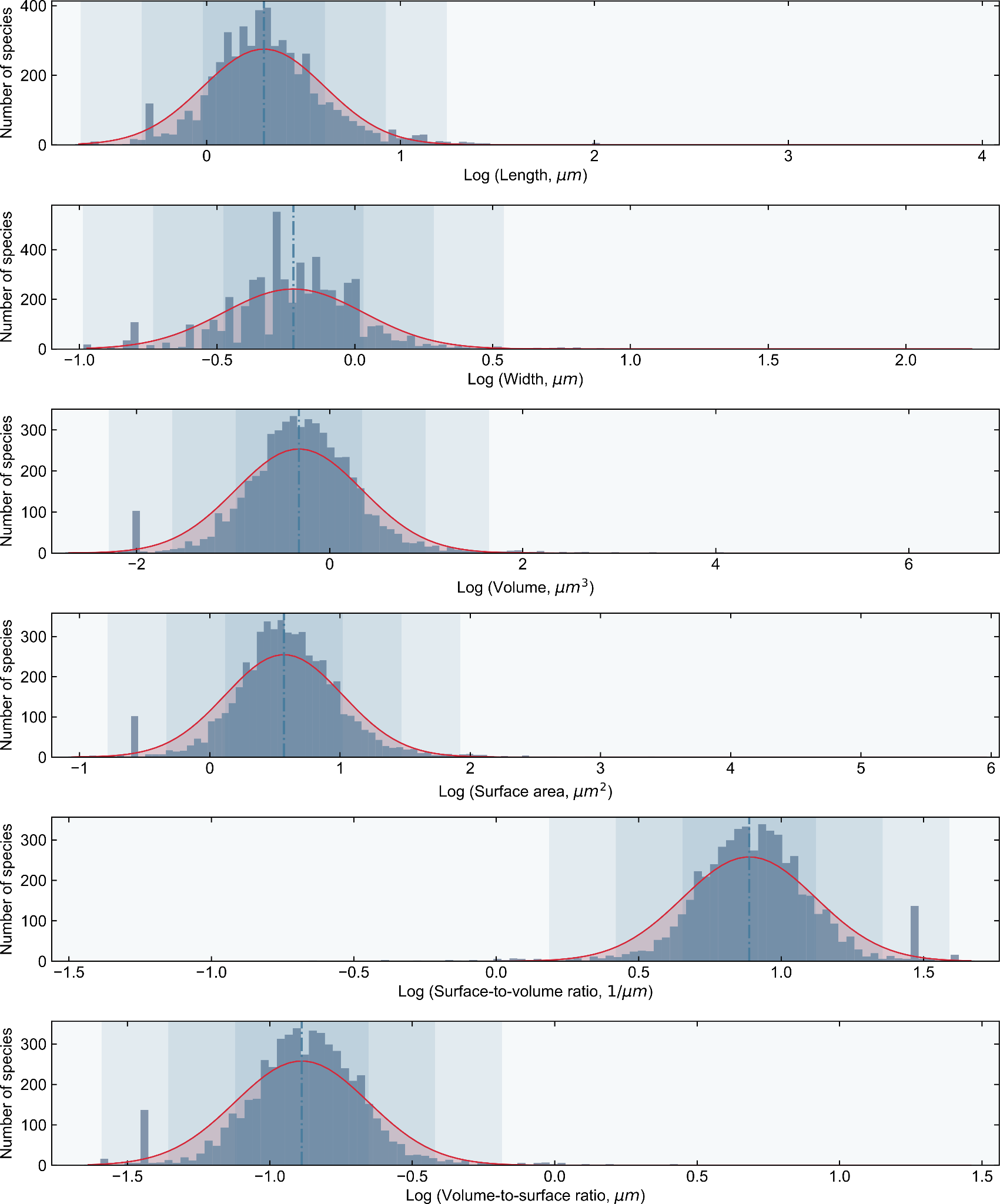
**Figure S10. Related to section “A multiplicity of drivers”.** Distributions of cell size according to different measures. In gray is a histogram of binned data. Red solid lines represent the log-normal distribution fitted to the data. The dashdot line indicates the mean. One, two, and three standard deviations from the mean are shown as a blue gradient. Notice that the artifactual peak is due to rounding effects across derived metrics (*V, SA, S/V, V/S*).

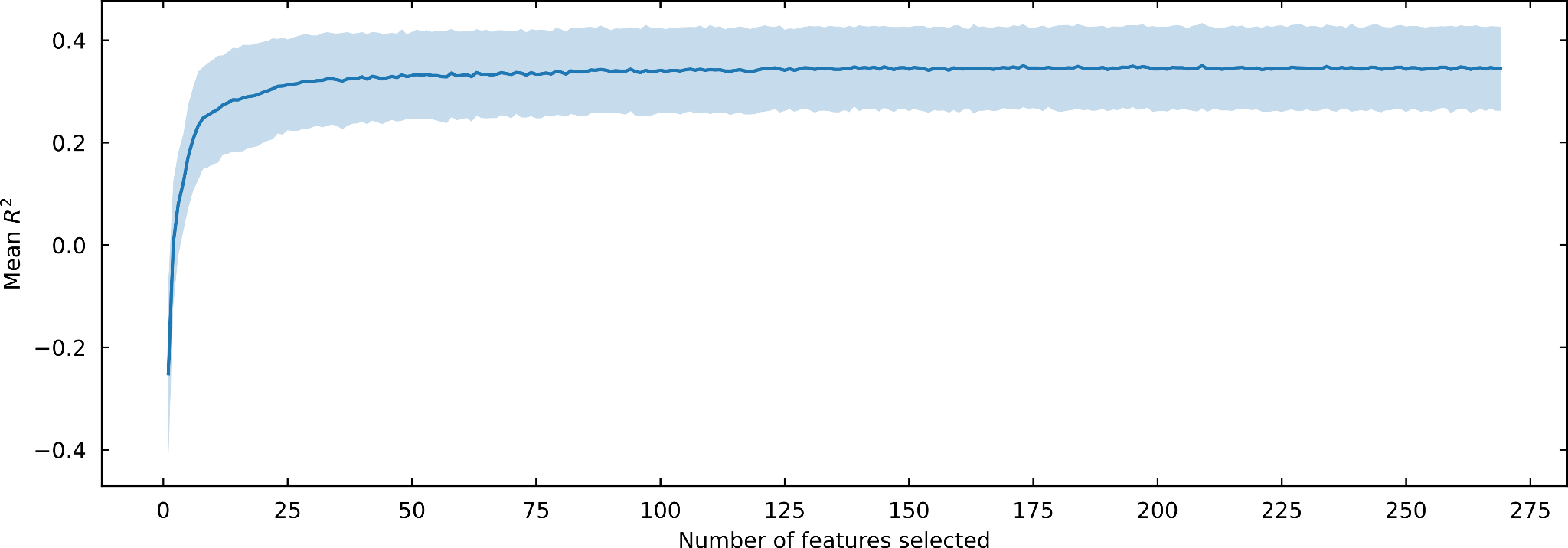

**Figure S11**. **Related to section “A multiplicity of drivers”.** Saturation curve of the number of features for cell size prediction. A total of 269 genomic features were iteratively reduced through recursive feature elimination with cross-validation (RFECV) with a random forest regression model to predict cell size. In each iteration, the retained features were evaluated using the coefficient of determination (*R*^2^) calculated from 10 repeats of 5-fold cross-validation (*y*-axis). Mean and standard deviation of the result are indicated by the line and the band, respectively.

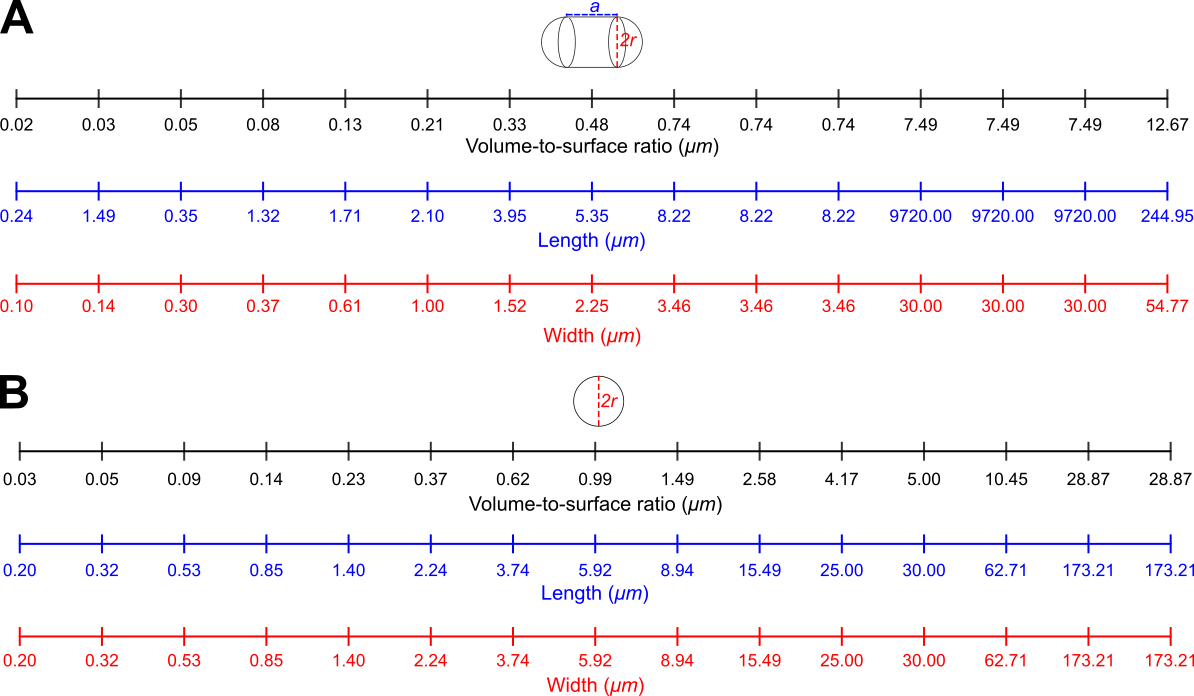

**Figure S12. Related to section “Data collection and curation”.** Mapping between linear measures of cell size for rod (A) and spherical (B) cells. A capsule and sphere are shown to demonstrate the calculation of volume-to-surface ratio (see Methods). a = length - width and r = width/2.

#### **Supplemental tables**

**Table S1. Related to Figure 1.** Comparing medians across cell shapes using the Mann-Whitney *U* test. *U*: values of the Mann-Whitney *U* statistical test and *P*: *p*-value.

| **Shape pair** | ***U*** | ***P ^a^*** |
| --- | --- | --- |
| **Rods-Cocci** | 0.2 | <0.001 |
| **Rods-Spirals** | 0.48 | <0.001 |
| **Rods-Filaments** | 0.54 | <0.001 |
| **Cocci-Spirals** | 0.5 | <0.001 |
| **Cocci-Filaments** | 0.47 | <0.001 |
| **Spirals-Filaments** | 0.18 | 0.52 |
| ^a^ *P*-values were corrected using the Benjamini-Hochberg method. | | |

**Table S2. Related to Figure 2B.** Comparing volume-to-surface ratio across phyla with the Mann-Whitney *U* test. *U*: test statistic, *P*: *p*-value.

| **Phylum pair** | ***U*** | ***P ^a^*** |
| --- | --- | --- |
| **Proteobacteria-Bacteroidetes** | 741085.5 | <0.001 |
| **Proteobacteria-Firmicutes** | 687059.0 | <0.001 |
| **Proteobacteria-Actinobacteria** | 626973.5 | <0.001 |
| **Proteobacteria-Tenericutes** | 6782.5 | <0.001 |
| **Proteobacteria-Euryarchaeota** | 105942.5 | <0.001 |
| **Proteobacteria-Spirochaetes** | 23225.0 | <0.001 |
| **Proteobacteria-Cyanobacteria** | 7550.5 | <0.001 |
| **Bacteroidetes-Firmicutes** | 231321.5 | <0.001 |
| **Bacteroidetes-Actinobacteria** | 340779.0 | <0.05 |
| **Bacteroidetes-Tenericutes** | 5243.5 | <0.001 |
| **Bacteroidetes-Euryarchaeota** | 37085.5 | <0.001 |
| **Bacteroidetes-Spirochaetes** | 17589.5 | <0.001 |
| **Bacteroidetes-Cyanobacteria** | 2066.0 | <0.001 |
| **Firmicutes-Actinobacteria** | 198319.0 | <0.001 |
| **Firmicutes-Tenericutes** | 1525.5 | <0.001 |
| **Firmicutes-Euryarchaeota** | 48497.0 | <0.05 |
| **Firmicutes-Spirochaetes** | 6298.0 | <0.001 |
| **Firmicutes-Cyanobacteria** | 3648.0 | <0.001 |
| **Actinobacteria-Tenericutes** | 3810.5 | <0.001 |
| **Actinobacteria-Euryarchaeota** | 31379.5 | <0.001 |
| **Actinobacteria-Spirochaetes** | 12235.0 | <0.001 |
| **Actinobacteria-Cyanobacteria** | 2034.0 | <0.001 |
| **Tenericutes-Euryarchaeota** | 786.5 | <0.001 |
| **Tenericutes-Spirochaetes** | 4914.0 | <0.001 |
| **Tenericutes-Cyanobacteria** | 4.5 | <0.001 |
| **Euryarchaeota-Spirochaetes** | 1333.0 | <0.001 |
| **Euryarchaeota-Cyanobacteria** | 801.5 | <0.001 |
| **Spirochaetes-Cyanobacteria** | 51.0 | <0.001 |
| ^a^ P-values were corrected using Benjamini-Hochberg correction. | | |

**Table S3. Related to Figure 2B.** Assessing kurtosis, skewness, and normality of distributions among phyla with the Kolmogorov-Smirnov test for goodness of fit. *μ*: mean, *σ*: standard deviation, *D*: value of the test statistic, *P*: *p*-value, *ɣ*: skewness, and K: kurtosis.

| **Phylum** | ***μ* ^a^** | ***σ* ^a^** | ***D*** | ***P* ^b^** | ***ɣ^c^*** | **K^d^** |
| --- | --- | --- | --- | --- | --- | --- |
| **Tenericutes** | 0.04 | 1.38 | 0.44 | <0.001 | 1.60 | 1.24 |
| **Spirochaetes** | 0.06 | 1.65 | 0.12 | 0.11 | -0.31 | -0.05 |
| **Bacteroidetes** | 0.12 | 1.46 | 0.03 | 0.33 | 0.16 | 0.29 |
| **Actinobacteria** | 0.12 | 1.46 | 0.03 | 0.35 | 0.75 | 4.33 |
| **Proteobacteria** | 0.14 | 1.52 | 0.04 | <0.05 | 1.40 | 17.31 |
| **Firmicutes** | 0.15 | 1.49 | 0.06 | <0.05 | 1.50 | 18.36 |
| **Euryarchaeota** | 0.15 | 1.61 | 0.09 | 0.33 | -0.80 | 1.29 |
| **Cyanobacteria** | 0.66 | 2.40 | 0.09 | 0.35 | 0.49 | 0.38 |
| ^a^ Values are reported in linear scale.  ^b^ *P* corrected using Benjamini-Hochberg correction.  ^c^ɣ < 0: left skewness; ɣ = 0: no skewness; ɣ > 0 right skewness.  ^d^K < 0: platykurtic; K = 0: mesokurtic; K > 0: leptokurtic. | | | | | | |

**Table S4. Related to Figure 4.** Phylogenetic signal across taxonomic groups. λ: value of Pagel’s λ, *σ*^2^: variance, Lnlik: log-likelihood, and *P: p*-value. Available as an Excel spreadsheet.

**Table S5. Related to section “Evolutionary models speak for the existence of an optimal cell size in prokaryotes”.** Summary of evolutionary models for cell size on a phylogenetic tree, without rescaling. OU model shows the highest likelihood and the smallest AIC. LnLik: log-likelihood, AIC: Akaike Information Criterion, *σ*^2^: rate of evolution of trait, *Z*_0_: trait value assigned at the root of the phylogenetic tree, *a:* pattern of rate decline, *α*: attractor strength, *θ:* optimum value.

| **Model** | **Parameters** | **Lnlik** | **AIC** | **Akaike weight** |
| --- | --- | --- | --- | --- |
| **Brownian motion** | σ^2^ = 0.17  Z_0_ = -0.89 | 1643.0 | -3282.0 | 0 |
| **Early-Burst** | a = -1.e-06  σ^2^ = 0.17  Z_0_ = -0.89 | 1643.0 | -3280.0 | 0 |
| **White-noise** | σ^2^ = 0.05  Z_0_ = -0.89 | 174.47 | -344.94 | 0 |
| **Ornstein-Uhlenbeck^*^** | **α = 1.95**  **σ^2^ = 0.23**  ***θ* = -0.93** | **1794.45** | **-3582.89** | **1** |
| **^*^**Best model according to Lnlik, AIC, and Akaike weights. | | | | |

**Table S6. Related to sections “What forces may be behind a prokaryotic cell size optimum?”, “Could S/V ratios prevent bacteria from evolving cells much larger than the norm?” and Figure S6.** Estimates of ordinary least squares regression between genome properties and volume-to-surface ratio. *ɑ*: Intercept, *β*: slope, *P*: *p*-value, *R*^2^: coefficient of determination, and AIC: Akaike Information Criterion value.

| **Predictor variable** | ***ɑ*** | **Standard error** | ***P_ɑ_*** | ***β*** | **Standard error** | ***P_β_*** | **R^2^** | **AIC** |
| --- | --- | --- | --- | --- | --- | --- | --- | --- |
| **Log genome size** | 6.75 | 0.02 | 0.00 | 0.21 | 0.02 | 0.00 | 0.07 | -866.5 |
| **Log number of proteins** | 3.68 | 0.02 | 0.00 | 0.18 | 0.02 | 0.00 | 0.06 | -1085 |
| **Arcsine**  **GC content** | 0.82 | 0.01 | 0.00 | 0.03 | 0.02 | 0.06 | 0.002 | -1674 |
| **Arcsine Coding density** | 1.19 | 0.01 | 0.00 | -0.04 | 0.01 | 0.00 | 0.02 | -4058 |
| **16S rRNA copies** | 2.58 | 0.19 | 0.00 | 0.85 | 0.21 | 0.00 | 0.01 | 5478 |
| **ΔENC’** | 0.17 | 0.01 | 0.00 | 0.02 | 0.01 | 0.008 | 0.004 | -2999 |

**Table S7. Related to section “What forces may be behind a prokaryotic cell size optimum?” and Figure S6.** Estimates of phylogenetic least squares regression between genome properties and volume-to-surface ratio. *ɑ*: Intercept, *β*: slope, *P*: *p*-value, and *R*^2^: coefficient of determination.

| **Predictor variable** | ***ɑ*** | **Standard error** | ***P_ɑ_*** | ***β*** | **Standard error** | ***P_β_*** | **R^2^** |
| --- | --- | --- | --- | --- | --- | --- | --- |
| **Log genome size** | 6.46 | 0.07 | <0.0001 | 0.007 | 0.01 | <0.0001 | 0.02 |
| **Log number of proteins** | 3.45 | 0.07 | <0.0001 | 0.07 | 0.01 | <0.0001 | 0.02 |
| **Arcsine**  **GC content** | 0.75 | 0.04 | <0.0001 | -0.004 | 0.007 | 0.6 | -0.0005 |
| **Arcsine Coding density** | 1.23 | 0.03 | <0.0001 | -0.008 | 0.005 | 0.12 | 0.001 |
| **16S rRNA copies** | 1.61 | 1,65 | 0.33 | -0.22 | 0.32 | 0.49 | -0.0004 |
| **ΔENC’** | 0.09 | 0.03 | < 0.05 | 0.006 | 0.006 | 0.38 | -0.0002 |

**Table S8. Related to sections “What forces may be behind a prokaryotic cell size optimum?”, “Usefulness and limitations of the dataset”, and Figure S7.** Estimates of ordinary least squares regression between genome properties and volume. *ɑ*: Intercept, *β*: slope, *P*: *p*-value, *R*^2^: coefficient of determination, and AIC: Akaike Information Criterion value.

| **Predictor variable** | ***ɑ*** | **Standard error** | ***P_ɑ_*** | ***β*** | **Standard error** | ***P_β_*** | **R^2^** | **AIC** |
| --- | --- | --- | --- | --- | --- | --- | --- | --- |
| **Log genome size** | 6.57 | 0.01 | 0.00 | 0.06 | 0.01 | 0.00 | 0.05 | -838.7 |
| **Log number of proteins** | 3.54 | 0.01 | 0.00 | 0.06 | 0.01 | 0.00 | 0.05 | -1061 |
| **Arcsine**  **GC content** | 0.79 | 0.01 | 0.00 | -0.02 | 0.01 | <0.05 | 0.005 | -1679 |
| **Arcsine Coding density** | 1.22 | 0.01 | 0.00 | -0.01 | 0.01 | 0.00 | 0.03 | -4064 |
| **16S rRNA copies** | 1.91 | 0.05 | 0.00 | 0.32 | 0.08 | 0.00 | 0.01 | 5478 |
| **ΔENC’** | 0.15 | 0.01 | 0.00 | 0.003 | 0.003 | 0.32 | 0.00 | -2993 |

**Table S9. Related to sections “What forces may be behind a prokaryotic cell size optimum?” and Figure S7.** Estimates of phylogenetic least squares regression between genome properties and volume. *ɑ*: Intercept, *β*: slope, *P*: *p*-value, and *R*^2^: coefficient of determination.

| **Predictor variable** | ***ɑ*** | **Standard error** | ***P_ɑ_*** | ***β*** | **Standard error** | ***P_β_*** | **R^2^** |
| --- | --- | --- | --- | --- | --- | --- | --- |
| **Log genome size** | 6.41 | 0.07 | <0.0001 | 0.03 | 0.01 | <0.0001 | 0.02 |
| **Log number of proteins** | 3.4 | 0.07 | <0.0001 | 0.02 | 0.01 | <0.0001 | 0.02 |
| **Arcsine**  **GC content** | 0.75 | 0.04 | <0.0001 | -0.001 | 0.002 | 0.64 | -0.0006 |
| **Arcsine Coding density** | 1.24 | 0.03 | <0.0001 | -0.004 | 0.002 | <0.05 | 0.003 |
| **16S rRNA copies** | 1.79 | 1/63 | 0.27 | -0.05 | 0.11 | 0.66 | -0.0006 |
| **ΔENC’** | 0.08 | 0.03 | <0.05 | 0.0009 | 0.002 | 0.67 | -0.0006 |

**Table S10. Related to section “What forces may be behind a prokaryotic cell size optimum? “** Estimates of ordinary least squares regression between KEGG pathways and volume-to-surface ratio. The fraction of total number of base pairs of all genes in a pathway divided by genome size was used as a measure of ecological/evolutionary relevance of a pathway. *ɑ*: Intercept, *β*: slope, *P*: *p*-value, *R*^2^: coefficient of determination, and AIC: Akaike Information Criterion value. Available as an Excel spreadsheet.

**Table S11. Related to section “A multiplicity of drivers” and Figure S10.** Assessing kurtosis, skewness, and normality of cell size distributions according to different metrics with the Kolmogorov-Smirnov test for goodness of fit. *μ*: mean, *σ*: standard deviation, *D*: value of the test statistic, *P*: *p*-value, and K: kurtosis.

| **Metric** | ***μ*^a^** | ***σ*^a^** | ***D*** | ***P*^b^** | ***ɣ^c^*** | **K^d^** |
| --- | --- | --- | --- | --- | --- | --- |
| **Log length** | 1.97 | 2.06 | 0.08 | <0.001 | 1.09 | 6.13 |
| **Log width** | 0.60 | 1.80 | 0.08 | <0.001 | 1.01 | 7.17 |
| **Log volume** | 0.48 | 4.53 | 0.06 | <0.001 | 1.15 | 10.38 |
| **Log surface area** | 3.71 | 2.83 | 0.06 | <0.001 | 1.09 | 10.03 |
| **Log surface-to-volume ratio** | 7.73 | 1.71 | 0.06 | <0.001 | -0.95 | 7.95 |
| **Log volume-to-surface ratio** | 0.13 | 1.71 | 0.06 | <0.001 | 0.95 | 7.95 |
| ^a^Values are reported in linear scale.  ^b^P-values corrected using Benjamini-Hochberg correction.  ^c^ɣ < 0: left skewness; ɣ = 0: no skewness; ɣ > 0 right skewness.  ^d^K < 0: platykurtic; K = 0: mesokurtic; K > 0: leptokurtic. | | | | | | |
